# Direct phosphorylation of HY5 by SPA1 kinase to regulate photomorphogenesis in Arabidopsis

**DOI:** 10.1101/2020.09.10.291773

**Authors:** Wenli Wang, Inyup Paik, Junghyun Kim, Xilin Hou, Sibum Sung, Enamul Huq

## Abstract

ELONGATED HYPOCOTYL5 (HY5) is a key transcription factor which promotes photomorphogenesis by regulating complex downstream growth programs. Previous studies suggest that the regulation of HY5 mainly depends on the CONSTITUTIVE PHOTOMORPHOGENIC1 (COP1) - SUPPRESSOR OF PHYTOCHROME A-105 (SPA) E3 ubiquitin ligase complex, which degrades positively acting transcription factors of light signaling to repress photomorphogenesis in the dark. SPA proteins function not only as a component of the E3 ubiquitin ligase complex but also as a kinase of PHYTOCHROME INTERACTING FACTOR1 (PIF1) through its N-terminal kinase domain. Here, we show that HY5 is a new substrate of SPA1 kinase. SPA1 can directly phosphorylate HY5 *in vitro* and *in vivo*. We also demonstrate that unphosphorylated HY5 strongly interacts with both COP1 and SPA1 than phosphorylated HY5, is the preferred substrate for degradation, whereas phosphorylated HY5 is more stable in the dark. In addition, unphosphorylated HY5 actively binds to the target promoters, and is physiologically more active form. Consistently, the transgenic plants expressing unphosphorylated mutant of HY5 displays enhanced photomorphogenesis. Collectively, our study revealed that SPA1 fine-tunes the stability and the activity of HY5 to regulate photomorphogenesis.

## INTRODUCTION

Light is a key environmental factor that influences diverse developmental processes throughout the entire plant life cycle (Whitelam and Halliday, 2007). Plants have evolved four classes of photoreceptors to monitor the surrounding light conditions: red (R)/far-red (FR) light–sensing phytochromes, blue/UV-A light–sensing cryptochromes and phototropins, and UV-B light– sensing UVR8 (Chen *et al*., 2004, Paik and Huq, 2019). Interestingly, all the light signals perceived by different photoreceptors converge to a downstream transcription factor ELONGATED HYPOCOTYL5 (HY5) to control diverse growth programs (Gangappa and Botto, 2016). For example, dark-grown Arabidopsis seedlings undergo skotomorphogenesis, displaying closed, yellowish cotyledons and long hypocotyls. Upon light irradiation, seedlings undergo photomorphogenesis which includes open, wide and green cotyledons and short hypocotyls (Gommers and Monte, 2018). The dark to light transition mainly causes the accumulation of HY5 proteins and then triggers cascades of downstream gene expressions. Indeed, HY5 regulates nearly one third of Arabidopsis genes (Lee *et al*., 2007, Gangappa and Botto, 2016), and a wide range of plant developmental programs, including flowering time, chlorophyll and anthocyanin biosynthesis, primary and lateral root development, shade and high temperature responses (Oyama *et al*., 1997, Ang *et al*., 1998, Holm *et al*., 2002, Andronis *et al*., 2008, Nozue *et al*., 2015, Gangappa and Botto, 2016).

The level of HY5 protein is regulated by CONSTITUTIVE PHOTOMORPHOGENIC1 (COP1) - SUPPRESSOR OF PHYTOCHROME A-105 (SPA) E3 ubiquitin ligase complex (Hoecker, 2017). Both COP1 and SPA are crucial repressors of photomorphogenesis. COP1 protein is enriched in nucleus in the dark and depleted from nucleus in the light. Thus, in darkness, COP1-SPA complex induces ubiquitination and degradation of HY5 and possibly other positive transcription factors in the nucleus (Hoecker, 2017, Han *et al*., 2020). However, upon light irradiation, the activity of COP1-SPA E3 ubiquitin ligase complex is inhibited by photoreceptors (Ordoñez-Herrera *et al*., 2015, Sheerin *et al*., 2015, Xu *et al*., 2015). COP1 is also excluded from the nucleus in response to light (Subramanian *et al*., 2004, Pacín *et al*., 2014, Balcerowicz *et al*., 2017). The reduction of COP1 in the nucleus and the light-induced inhibition of COP1 activity contribute to the accumulation of HY5 and other positive regulators to promote photomorphogenesis.

The function of COP1 as a RING-type E3 ubiquitin ligase is evolutionarily conserved in higher eukaryotes. It consists of an N-terminal zinc finger, a central coiled-coil (CC), and a C-terminal WD40 repeats domain, which is essential for proper COP1 function (Deng *et al*., 1992, Han *et al*., 2020). SPA family of genes are only found in the green lineages. Arabidopsis has four *SPA* genes (*SPA1*-*SPA4*) (Laubinger *et al*., 2004, Hoecker, 2017). SPA family of proteins also contain the central CC and the C-terminal WD40 repeats domain, which function similar to the respective domains of COP1 (Hoecker *et al*., 1999). In addition, SPA proteins contain an N-terminal Ser/Thr kinase domain which was recently found to have kinase activity on PHYTOCHROME INTERACTING FACTOR1 (PIF1), a negative regulator in photomorphogenesis (Paik *et al*., 2019). SPA1 and COP1 can interact with each other through their CC domains. SPA proteins are important for COP1 function, the presence of SPA proteins can enhance the activity of COP1 *in vivo* (Saijo *et al*., 2003, Seo *et al*., 2003). Both *cop1* and *spaQ* (*spa1spa2spa3spa4* quadruple) mutants exhibit constitutive photomorphogenesis in the darkness. In plants, COP1 forms multiple complexes with SPA proteins in a tissue- and developmental stage-specific manner. COP1-SPA complex promotes ubiquitination and subsequent degradation of multiple transcription factors (Hoecker, 2017).

Previous studies showed that HY5 interacts with the WD40 repeat domain of both COP1 and SPA1 through its N-terminal domain (Torii *et al*., 1998, Hardtke *et al*., 2000, Hoecker, 2017). Given the known physical interactions and SPA kinase activity, it prompted us to examine whether HY5 is a new substrate of SPA kinase. Here, we provide evidence that HY5 is a new target of SPA kinase. SPA1 directly phosphorylates HY5 proteins both *in vitro* and *in vivo*. We found that phosphorylated and unphosphorylated HY5 proteins have different binding affinity to COP1-SPA complex, which results in different degradation rate of HY5 proteins in the dark. In addition, the phosphorylation of HY5 proteins also modulates its activity in light response. Thus, our study shows that the phosphorylation of HY5 by SPA is necessary for fine tuning photomorphogenesis.

## RESULTS

### SPA1 can directly phosphorylate Serine-36 on HY5 proteins *in vitro*

Our recent study shows SPA1 acts as a Ser/Thr kinase of PIF1 (Paik *et al*., 2019). Since COP1-SPA E3 ubiquitin ligase complex interacts with HY5 proteins in the dark to promote its degradation, we hypothesize that HY5 might be a new substrate of SPA kinase. To test whether SPA can phosphorylate HY5 proteins, we first purified strep-tagged full-length SPA1 protein from a eukaryotic expression host (*Pichia pastoris*) and GST-tagged HY5 from bacteria (*E. coli*), and performed *in vitro* kinase assay. We found that SPA1 directly phosphorylates HY5 proteins *in vitro* in a concentration-dependent manner (Figure 1A). To check if this phosphorylation activity is changed by the incubation time, we conducted a time gradient kinase assay (Figure 1B). With increasing HY5 proteins and incubation time, we observed stronger phosphorylation signals. These results suggest that SPA1 phosphorylates HY5 in a concentration- and time-dependent manner (Figure 1A, B).

**Figure 1.**
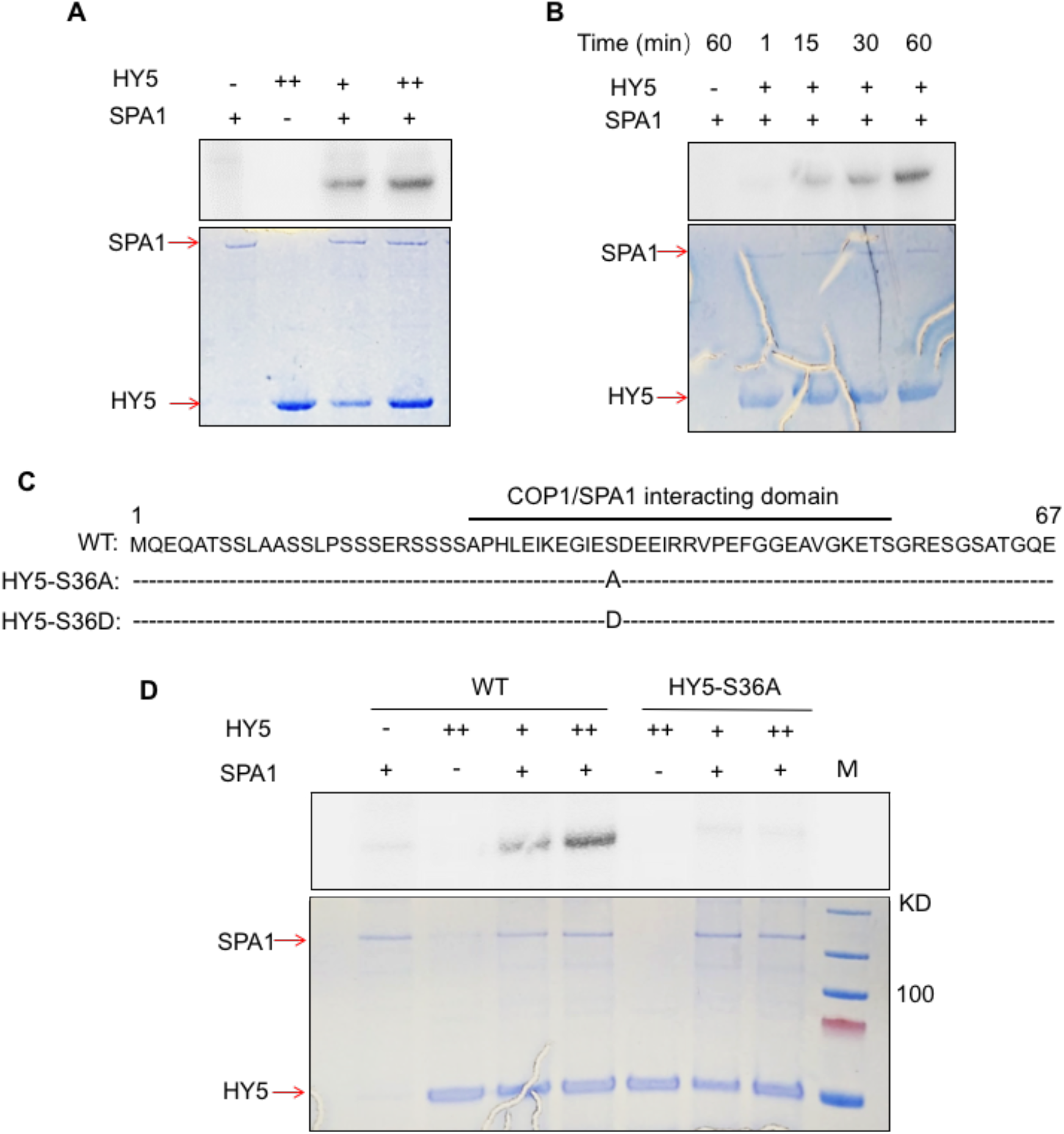
SPA1 acts as a Ser/Thr protein kinase and directly phosphorylates HY5 on Serine-36 *in vitro*. **(A)** Full-length SPA1 protein purified from *Pichia pastoris* phosphorylates HY5 protein *in vitro* in a concentration-dependent manner (autoradiogram on top panel). The bottom panel shows the protein levels in a Coomassie-stained gel. **(B)** Time-dependent kinase assays of full-length SPA1 on HY5 (autoradiogram on top panel). Bottom panel shows the protein levels in a Coomassie-stained gel. **(C)** Single phosphorylation site of HY5 is located at the 36th Serine (Ser-36) in its COP1 and SPA1 interacting domain. Ser-36 was then replaced with alanine (S36A) and aspartic acid (S36D) to generate a non-phosphorylation form and a phospho-mimicking form of HY5 respectively. **(D)** SPA1 protein purified from *Pichia pastoris* phosphorylates wild type HY5 but not the non-phosphorylation mutant of HY5 *in vitro* (autoradiogram on top panel). The bottom panel shows the protein levels in a Coomassie- stained gel. M indicates a protein marker.

To map the phosphorylation sites in HY5, we used GST-HY5 to conduct *in vitro* phosphorylation assays using SPA1 as a kinase. Mass-spectrometry analysis revealed a single phosphorylation site (Ser-36), which is located at the 36th serine at the N-terminus of HY5. A previous study showed that a deletion of the first 40 amino acids of HY5 completely abolished the interaction with COP1 (Intro_35). A 36 amino acid stretch between 25th and 60th residues of HY5 proteins was then defined as COP1 interacting domain. In addition, HY5 interacts with SPA1 through its N-terminal domain. Thus, the mapped phosphorylation site of HY5 is within its interaction domain for both COP1 and SPA (Figure 1C). To address the significance of the phosphorylation site, we replaced the Ser-36 with alanine (S36A) or aspartic acid (S36D) to generate non-phosphorylation mutant and phospho-mimicking mutant, respectively. *In vitro* kinase assay showed mutant recombinant protein (HY5-S36A) cannot be phosphorylated by SPA1, supporting that Ser-36 residue is indeed the single phosphorylation site of HY5 (Figure 1D).

### SPAs are necessary for phosphorylation of HY5 *in vivo*

To investigate the phosphorylation status of HY5 *in vivo*, we generated HY5 overexpressing transgenic plants (HY5-GFP) in wild type (WT) background and purified HY5-GFP proteins from transgenic seedlings grown in the dark or in the light. In immunoblot analysis, we observed band mobility shift after treatment with the native calf intestinal phosphatase (CIP) in both dark and light grown seedlings (Figure 2A and Supplemental Figure 1A, B), indicating that HY5 is phosphorylated under both dark and light conditions.

**Figure 2.**
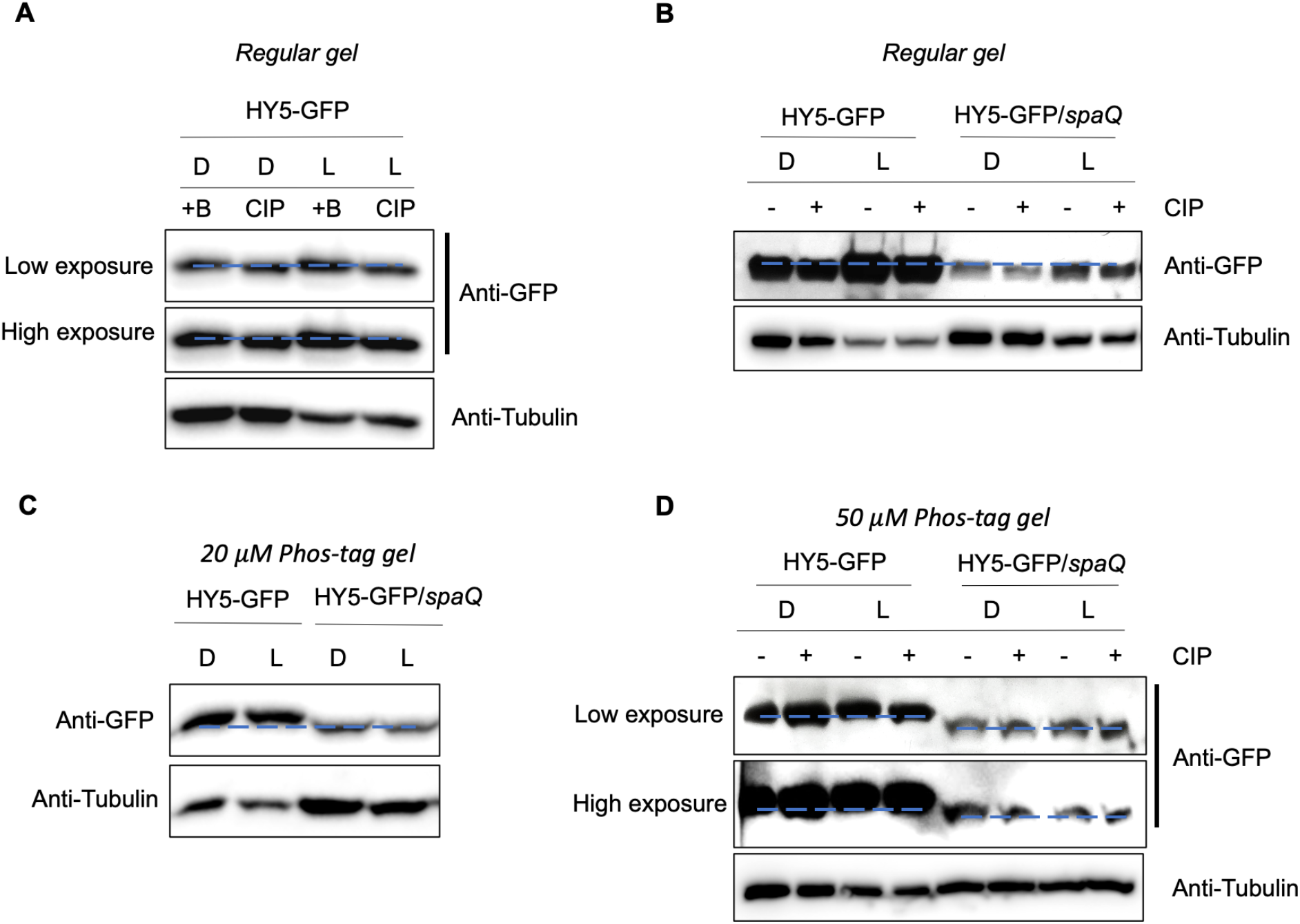
SPAs are necessary for the phosphorylation of HY5 *in vivo*. **(A)** Immunoblots showing phosphorylation of HY5-GFP under both dark and light conditions. Seedlings were grown in either dark (D) or continuous light (L) for 4 days before sampling for protein extraction. The slow-migrating bands are a phosphorylated form of HY5-GFP as indicated by the phosphatase treatment. CIP, Calf intestinal Alkaline Phosphatase; +B, inactivated boiled CIP. **(B)** Immunoblots showing a defect of HY5 phosphorylation in *spaQ* background compared to wild type. Seedlings were grown in either dark or continuous light for 4 days before sampling for protein extraction. The slow-migrating bands are phosphorylated forms of HY5-GFP as indicated by the phosphatase treatment. +, with CIP treatment; -, without CIP treatment. **(C and D)** Immunoblots showing a defect in phosphorylation of HY5-GFP in *spaQ* background compared to wild type in gels containing 20 **(C)** and 50 **(D)** µM phos-tag. The slow-migrating bands are phosphorylated forms of HY5-GFP. Tubulin proteins were used as loading control. +, with CIP treatment; -, without CIP treatment.

To examine the *in vivo* effect of SPAs on the HY5 phosphorylation, we then generated transgenic plants overexpressing HY5-GFP in *spaQ* background (HY5-GFP/*spaQ*). Strikingly, the phosphorylation and band shift of HY5 by CIP treatment observed in WT were completely abolished in the *spaQ* mutants both grown in the dark and light (Figure 2B). The HY5 phosphorylation status was further examined by utilizing Phos-tag containing SDS-PAGE gels (Figure 2C, D and Supplemental Figure 1C). In wild type plants, HY5-GFP showed clear mobility shift after CIP treatment in the presence of 20 µM Phos-tag (Supplemental Figure 1C). However, in *spaQ* mutant, no mobility shift was observed under these conditions and HY5-GFP showed a lower mobility shift compared to that in wild type (Figure 2C, D), suggesting a complete absence of phosphorylation of HY5 *in vivo*. Taken together, these results indicate that SPAs are responsible for HY5 phosphorylation *in vivo* under both light and dark conditions.

### SPA1 kinase domain is essential for its kinase activity and molecular function

SPA1 kinase domain is conserved in SPA1 sequences from multiple plants (Paik *et al*., 2019). The R517 residue in SPA1 is part of a conserved Glu-Arg salt bridge that defines eukaryotic protein kinases (Yang *et al*., 2012) and has recently been shown to be critical for its biological function in the dark (Holtkotte *et al*., 2016). To investigate the importance of SPA1-Ser/Thr kinase activity for HY5 phosphorylation, we used the point mutant version of SPA1 (mSPA1) that has the R517E mutation in the kinase domain. The *in vitro* kinase assay confirmed a significantly reduced phosphorylation activity of mSPA1 on HY5 (Supplemental Figure 2A).

To confirm the role of SPAs in the regulation of HY5, we measured the level of endogenous HY5 protein in different mutants and transgenic lines grown in the dark for 4 days (Supplemental Figure 3). As expected, we hardly detected HY5 protein signals in WT seedlings grown in the dark, whereas HY5 proteins were significantly accumulated in both *cop1* and *spaQ* mutants (Supplemental Figure 3A). In *spa* triple mutants (*spa123* and *spa124*), weaker bands of HY5 protein were detected, indicating that four SPA members have redundant functions in regulating HY levels (Supplemental Figure 3B). To clarify the biological role of SPA1 kinase activity, we then used the overexpressed SPA1 (LUC-SPA1) and mSPA1 (LUC-mSPA1) in *spa* quadruple mutants (*spaQ*) background. The HY5 level was more reduced by introduction of LUC-SPA1 compared to LUC-mSPA1 in the *spaQ* background (Supplemental Figure 2B), suggesting that the reduction in kinase activity of mSPA1 is deficient in degrading HY5 in the dark. This result is also consistent with the LUC-mSPA1 transgenic seedlings phenotype in the dark, which failed to rescue the constitutive photomorphogenesis, while LUC-SPA1 can largely rescue the constitutive photomorphogenic phenotype of *spaQ* (Supplemental Figure 2C).

**Figure 3.**
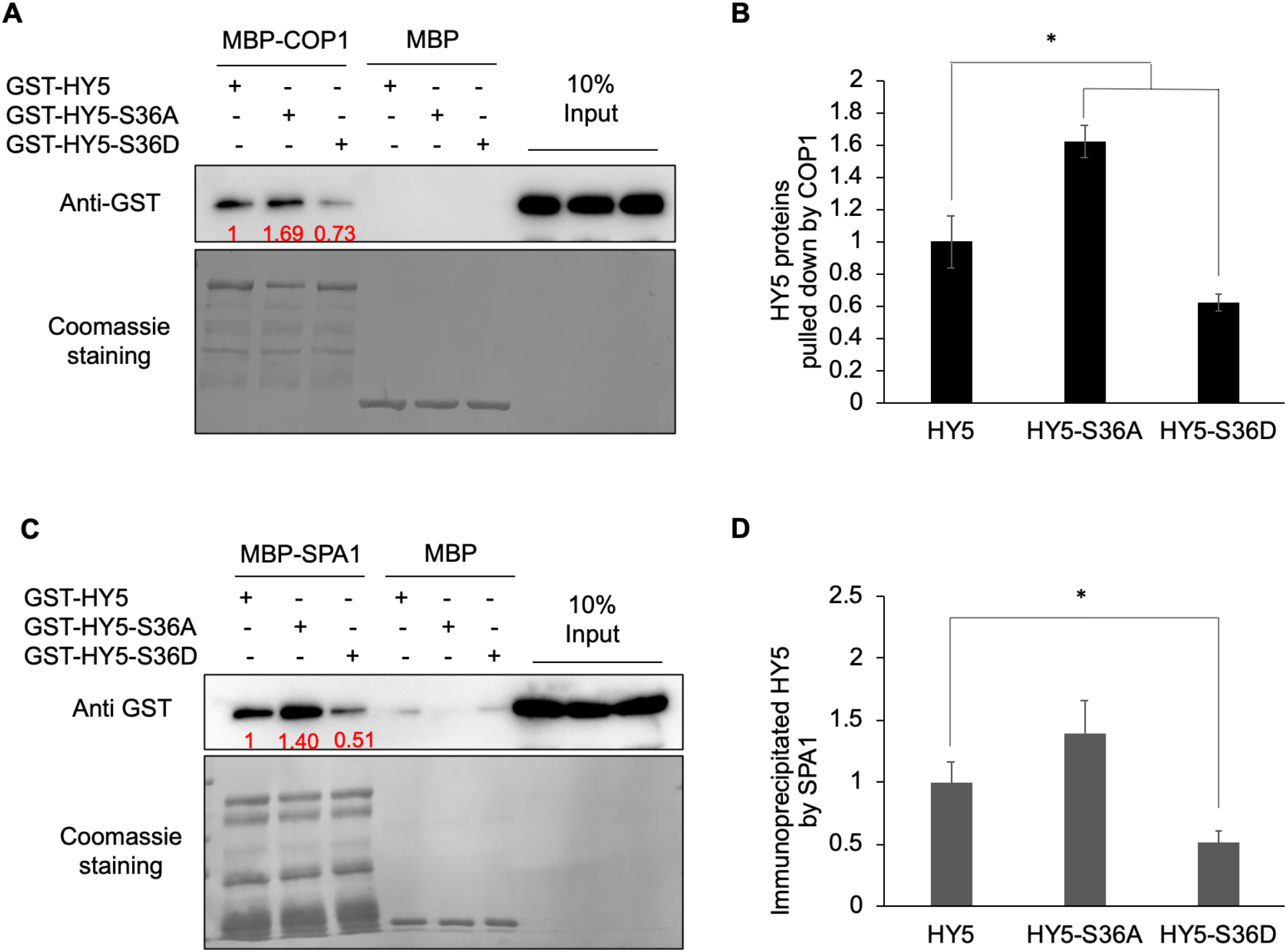
Phosphorylation of HY5 affects its interaction with COP1 and SPA1. **(A to D)**, Recombinant MBP-COP1, MBP-SPA1 and GST-HY5 (and mutant forms of HY5) proteins were purified from *Escherichia coli*. **(A)** *In vitro* pull-down assay shows that non-phosphorylated forms of HY5 (GST-HY5-S36A) has higher affinity to MBP-COP1. GST-HY5 and all other phosphorylation mutant proteins were pull-downed by MBP-COP1 using maltose agarose beads. The pellet fraction was eluted and analyzed by immunoblotting using anti-GST and anti-MBP antibodies or stained by Coomassie. **(B)** A bar graph showing the interaction between COP1 and wild type HY5, and mutant HY5 proteins. Error bars indicate SD (n = 3). The asterisk indicates a significant difference (P < 0.05). **(C)** *In vitro* pull-down assay shows that non-phosphorylated forms of HY5 (GST-HY5-S36A) has higher affinity to MBP-SPA1. GST-HY5 and all other phosphorylation mutant proteins were pull-downed by MBP-SPA1 using maltose agarose beads. The pellet fraction was eluted and analyzed by immunoblotting using anti-GST and anti-MBP antibodies or stained by Coomassie. **(D)** A bar graph showing the interaction between SPA1 and wild type HY5, and mutant HY5 proteins. Error bars indicate SD (n = 3). The asterisk indicates a significant difference (P < 0.05). The numbers below anti-GST blots (in **A** and **C**) indicate the relative band intensities of immunoprecipitated HY5 normalized to added COP1 and SPA1 proteins, respectively. The ratio of the first clear band was set to 1 for each blot.

### HY5 phosphorylation affects its interaction with COP1 and SPA1

Degradation of HY5 largely depends on its interaction with COP1-SPA complex in the nucleus (Intro_3&4). HY5 interacts with both COP1 and SPA1 through its N-terminal domain, and the phosphorylation site resides within the interacting domain (Figure 1C). We, therefore, examined whether the phosphorylation of HY5 affects its interactions with COP1 and SPA1.

To address this, we performed *in vitro* pull-down assays with purified fusion proteins, MBP-COP1 and MBP-SPA1. Each of the recombinant GST-fused wild-type HY5, HY5-S36A and HY5-S36D proteins was precipitated by MBP-COP1 or MBP-SPA1. Interestingly, the results show that non-phosphorylated form of HY5 (HY5-S36A) protein has higher affinity to both COP1 and SPA1, whereas phospho-mimicking form of HY5 (HY5-S36D) protein has significantly lower affinity to both COP1 and SPA1 compared with wild type HY5 (Figure 3). We also conducted *in vivo* co-IP assay with transgenic lines overexpressing similar amounts of HY5-GFP, HY5-S36A-GFP and HY5-S36D-GFP in *hy5* mutant background (Supplementary Figure 5A). When immunoprecipitated using GFP antibody, four times more COP1 protein were co-immunoprecipitated in HY5-S36A-GFP line than that in HY5-S36D-GFP line (Supplemental Figure 4). Taken together, both *in vitro* and *in vivo* data suggest that phosphorylation alters the affinity of HY5 to COP1-SPA complex and may also affect its accumulation and biological functions.

**Figure 4.**
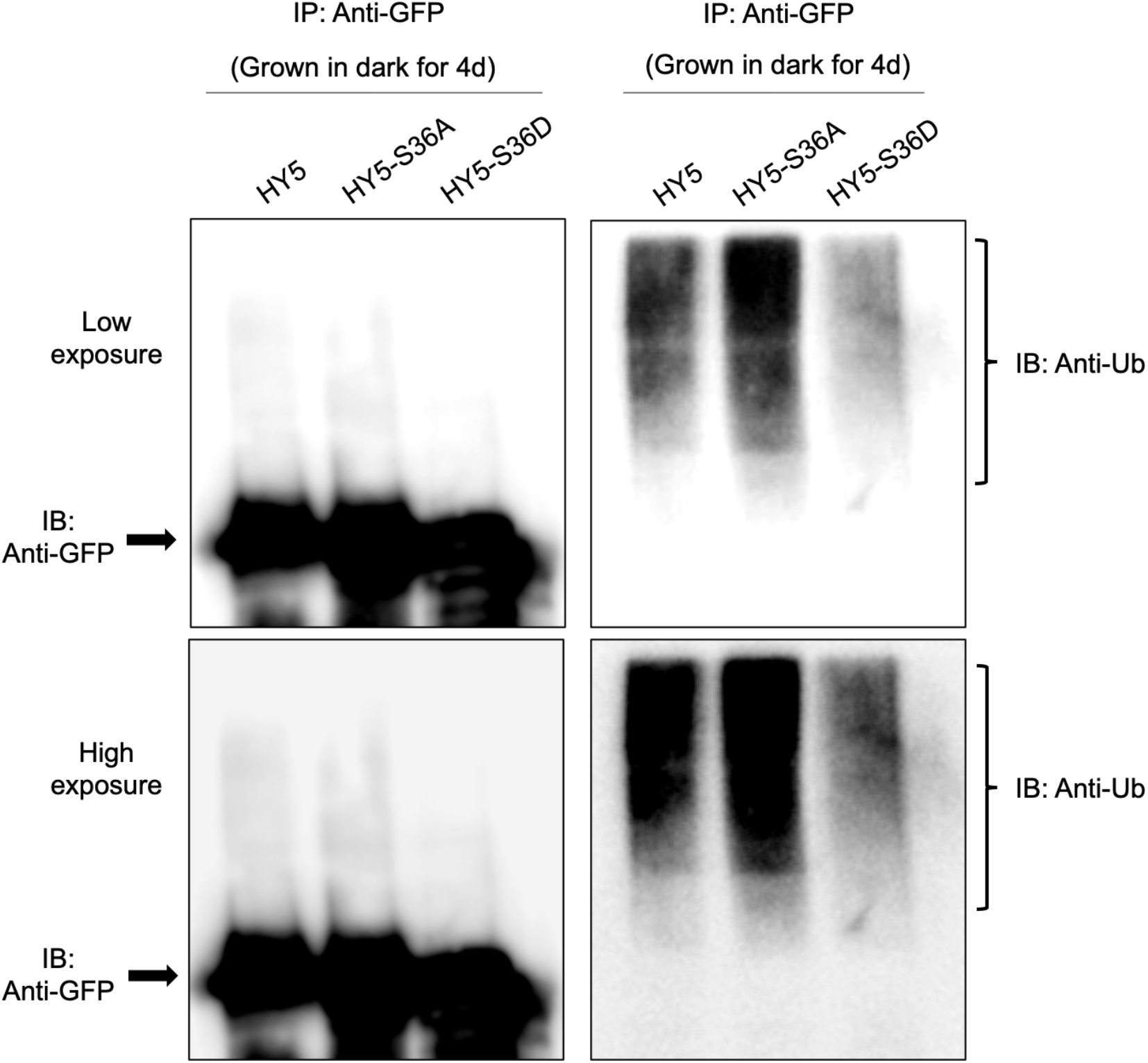
Phosphorylation alters HY5 ubiquitination level *in vivo*. Immunoblots showing the relative HY5 (-S36A/S36D)-GFP protein level (left) and their ubiquitination status in response to dark. Total protein was extracted from 4-day-old dark-grown seedlings pretreated with the proteasome inhibitor (40 µM bortezomib) for 4 h before protein extraction. HY5 (-S36A/S36D)-GFP was immunoprecipitated using anti-GFP antibody (rabbit) from protein extracts. The immunoprecipitated samples were then separated on 10% SDS-PAGE gels and probed with anti-GFP (left, Mouse) or anti-Ub (right) antibodies. The upper smear bands are polyubiquitinated HY5 (-S36A/S36D). Top: lower exposure; bottom: higher exposure. Arrows indicate the HY5 (-S36A/S36D)-GFP protein.

### Unphosphorylated HY5 degrades and accumulates faster than phosphorylated HY5

As a key positive regulator in seedlings photomorphogenesis, HY5 accumulates in response to the light and degrades in the dark (Osterlund *et al*., 2000). Previous studies have shown that the interaction between HY5 and COP1 or SPA1 is required for the degradation of HY5 (Osterlund *et al*., 2000, Saijo *et al*., 2003). HY5 is degraded in a polyubiquitination-dependent manner. As we observed that phosphorylation alters HY5-interaction affinity to COP1 and SPA1, we hypothesized that different interaction affinity may further affect HY5 stability. To verify our hypothesis, we examined the ubiquitination status of HY5 (-S36A/S-36D)-GFP *in vivo*. proteins were immunoprecipitated from HY5, HY5-S36A and HY5-S36D dark grown transgenic seedlings, pretreated with proteasome inhibitor (bortezomib), and then probed with anti-GFP and anti-Ub antibodies, respectively. Strikingly, the HY5-S36A has significantly higher ubiquitination level whereas HY5-S36D has significantly reduced ubiquitination level compared to wild type HY5 (Figure 4).

We also tested the protein levels of HY5, HY5-S36A and HY5-S36D transgenic seedlings which were grown in continuous light for four days and then transferred into dark for several hours (5, 10, 20 hrs) or grown in continuous dark for four days. The result shows that non-phosphorylated HY5 (HY5-S36A) is degraded faster during the light-to-dark transition, while phospho-mimicking forms of HY5 (HY5-S36D) is more stable, even in the dark (Figure 5A, B). Specifically, the degradation rate of non-phosphorylated HY5 was significantly increased after five hours of dark transition (Figure 5B). On the contrary, the degradation of phospho-mimicking HY5 was not obvious in the first ten hours of dark transition and its degradation was observed only after longer exposure of darkness (Figure 5B). Our results suggest that non-phosphorylated form of HY5 is the preferred substrate for degradation and phospho-mimicking forms of HY5 is not prone to be degraded. This is consistent with our observation that non-phosphorylated form of HY5 has higher binding affinity to COP1 and SPA1 and higher ubiquitination level, while phospho-mimicking forms of HY5 has much lower affinity to both proteins and lower ubiquitination level. These results indicate that the phosphorylation of HY5 results in weak interaction with COP1-SPA complex and subsequent ubiquitination status, making the HY5 proteins more stable in the dark.

**Figure 5.**
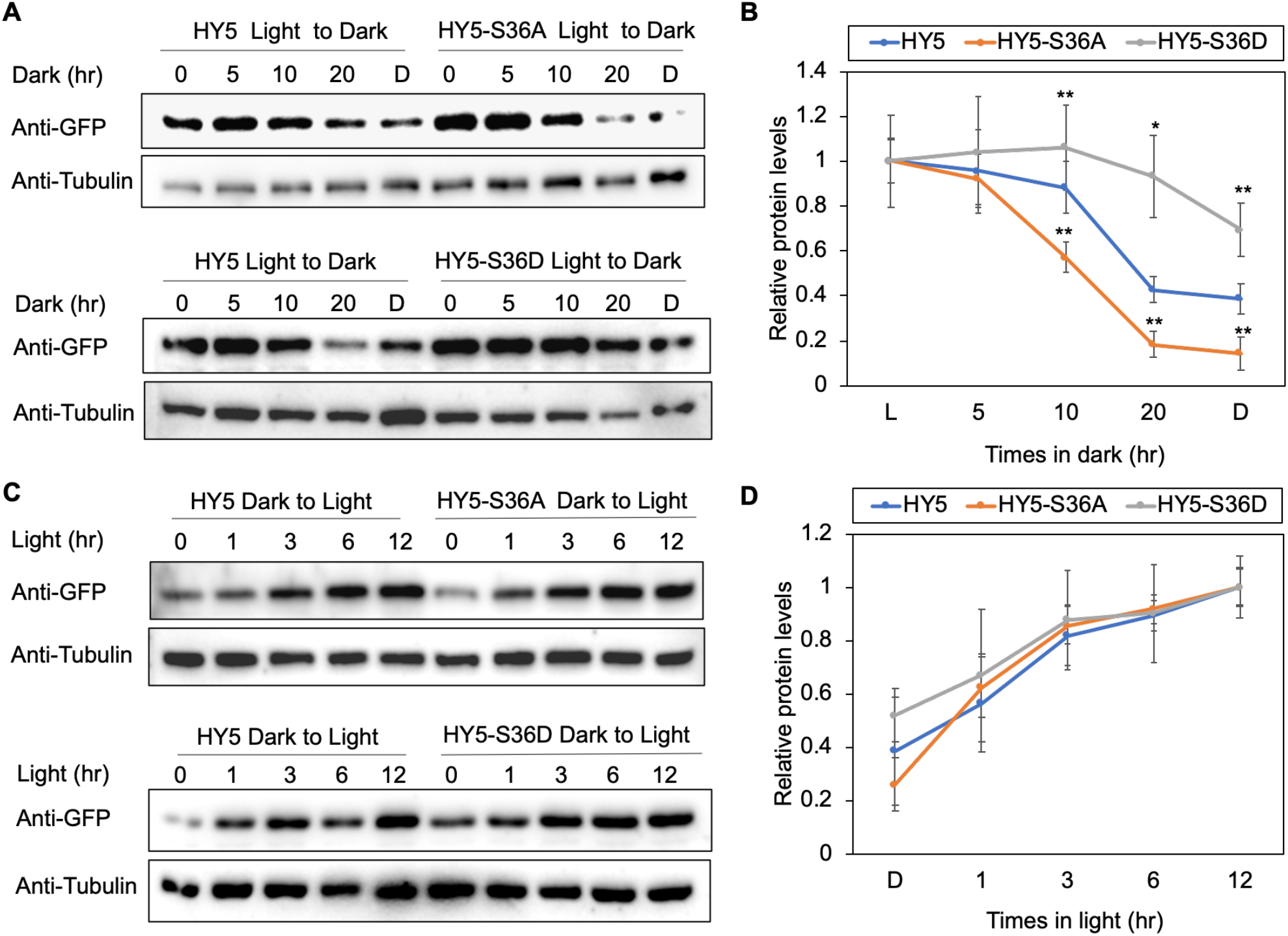
Phosphorylation alters HY5 degradation and accumulation rate. **(A)** Immunoblots showing the degradation pattern of HY5, HY5-S36A and HY5-S36D proteins in response to dark. Transgenic seedlings were grown in continuous light for 4 days and then transferred to Dark (D) for 5, 10 or 20 hrs, or grown in continuous dark for 4 days. Anti-GFP antibody and anti-tubulin antibody were used for this assay. Tubulin proteins were used as loading control. **(B)** A line graph shows the relative rate of degradation of HY5 in response to the dark treatment. The band intensities of HY5 and Tubulin were measured using the IMAGEJ tool. For each line, the HY5 level in the light (L) was set to 1 and the relative HY5 levels in response to dark were then calculated. Error bars indicate SD (n=3). The asterisk indicates a significant difference (P-value * <0.05, **<0.005). **(C)** Immunoblots showing the accumulation pattern of HY5, HY5-S36A and HY5-S36D proteins in response to light. Transgenic seedlings were grown in continuous dark for 4 days and then transferred to light for 1, 3, 6 or 12hrs. Anti-GFP antibody and anti-tubulin antibody were used for this assay. Tubulin proteins were used as loading control. **(D)** A line graph shows the relative accumulation rate of HY5 in response to the light treatment. The band intensities of HY5 and Tubulin were measured using the IMAGEJ tool. For each line, the HY5 level in the 12hrs light was set to 1 and the relative HY5 levels in response to light were then calculated. Error bars indicate SD (n=3).

In addition, we compared the accumulation rates of each mutant forms of HY5 during dark-to-light transition (Figure 5C, D). The accumulation of HY5 in response to light is very fast, and the accumulation occurs within one hour of light exposure. It reaches to the peak in three hours (Figure 5D). Non-phosphorylated form of HY5 appears to be more sensitive to the light irradiation, as it shows relatively rapid and higher HY5 protein accumulation within the first one hour after light irradiation, even though it is not statistically significant (Figure 5D). Taken together, our results suggest that phosphorylation of HY5 results in an altered response rate to dark and light. As a negative regulator of HY5, SPA1 phosphorylates HY5 and the phosphorylated HY5 remains stable in the dark, which suggests a COP1-SPA-HY5 negative feedback loop may exist.

### HY5 phosphorylation affects its physiological activity

Previous studies have reported that phosphorylation can affect binding affinity of HY5 to its target DNAs *in vitro* (Hardtke *et al*., 2000). To investigate whether phosphorylation alters the *in vivo* DNA bindings of HY5, we performed ChIP-qPCR assays. We selected several well-known HY5 target genes (*XTH15, EXP2, IAA19*, and *SAUR36*). Because HY5 protein accumulates to the peak after three hours of light activation during dark-to-light transition (Figure 5D), we performed ChIP-qPCR with transgenic seedlings grown in the dark for four days and then transferred to the light for additional three hours. Surprisingly, non-phosphorylated forms of HY5 enriched to the target loci significantly higher than the wild-type HY5. On the other hand, phospho-mimicking forms of HY5 enriched to target loci slightly lower than the wild-type HY5 (Figure 6A), indicating that unphosphorylated HY5 has higher affinity to its target DNA.

**Figure 6.**
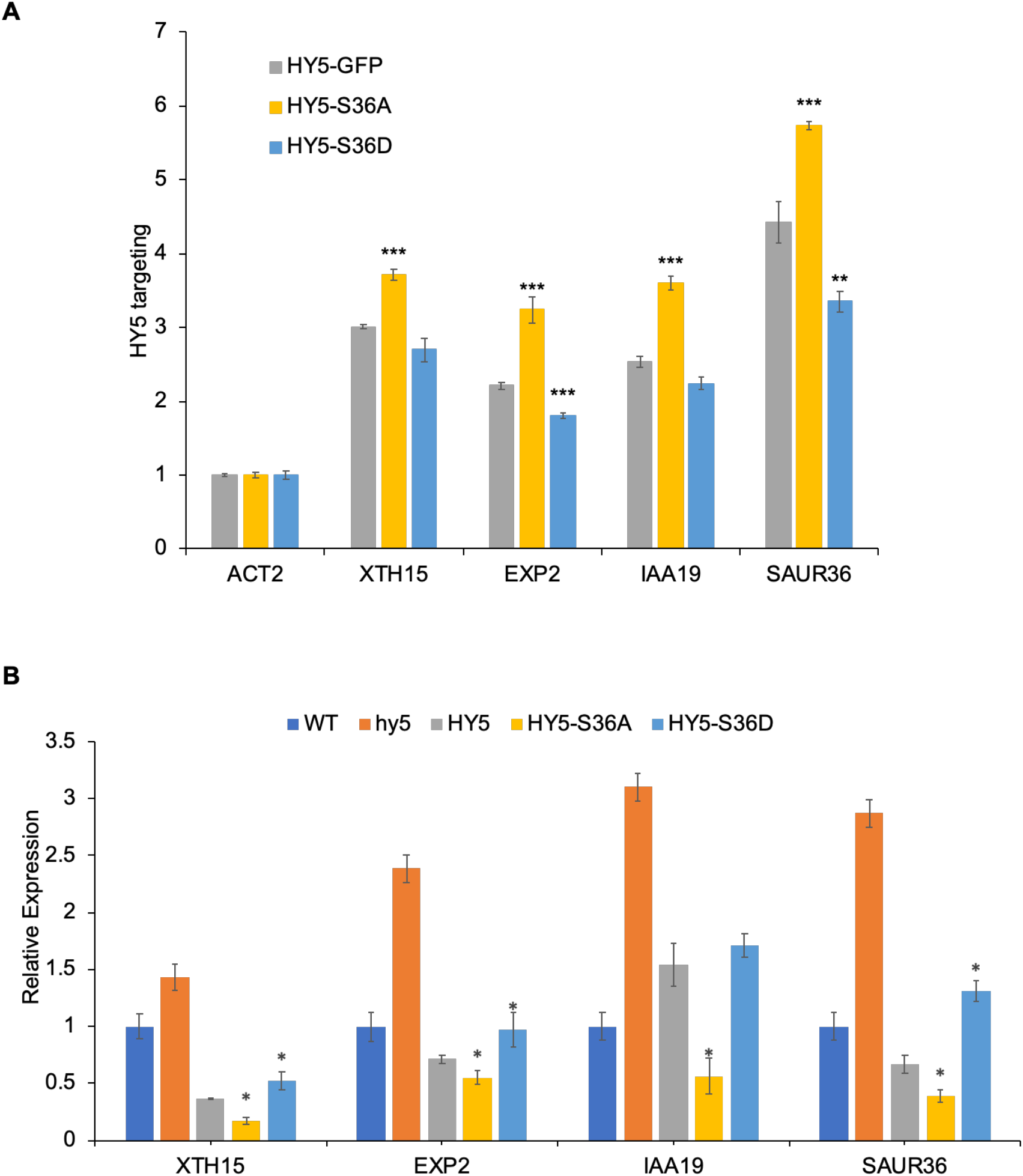
Unphosphorylated HY5 is physiologically more active in binding targets and regulating genes’ expression. **(A)** ChIP-qPCR results show the targeting of HY5 and two phosphorylation mutant proteins. For this assay, each transgenic seedlings were grown in dark for 4 days then transferred into light for additional 3hrs. The asterisk indicates a significant difference (P-value < 0.05, ** P-value < 0.005, *** P-value < 0.0005). Error bars indicate SD (n=3). **(B)** RT-qPCR results show transcript levels of selected HY5 target genes in HY5 and two phosphorylation mutant lines along with wild type and *hy5* as controls. For this assay, each transgenic seedlings were grown in dark for 4 days then transferred into light for additional 3hrs. The asterisk indicates a significant difference between HY5 and two mutant lines (P-value < 0.05). Error bars indicate SD (n = 3).

To examine whether the altered association of HY5 with its target loci affects their levels of transcript abundance, we then performed real-time qRT-PCR with the three transgenic lines grown under the same condition along with wild type and *hy5* as controls. Consistent with higher association, the expression of three auxin signaling pathway genes (*SAUR36, IAA19* and *EXP2*) and a growth gene (*XTH15*), which are known HY5 repressed genes, was strongly repressed in the HY5-S36A line (Figure 6B). However, HY5-S36D displayed higher expression of some of these target genes compared to the wild-type HY5. Our data suggested that the phosphorylation of HY5 affects the transcriptional activity of HY5 by changing its binding ability to target loci *in vivo*.

Furthermore, to check whether the phosphorylation mutants of HY5 result in any phenotypic changes, we measured the hypocotyl length of seedlings of the phosphorylation mutant lines grown under dark for 4 days or continuous red (Rc) or far-red (FRc) light for 4 days with increasing light intensities (Figure 7A, B). In both red light and far-red light conditions, *hy5* mutant showed relatively longer hypocotyl, which was rescued by overexpressing HY5 protein (Figure 7C, D). Notably, the HY5-S36A line showed shorter hypocotyls than others under most of Rc conditions and low intensity of FRc conditions (Figure 7A, B), even though all the lines had similar hypocotyl lengths in the dark. HY5-S36A hypocotyl length was strongly reduced once exposed to light. In addition, we also noticed that at the adult plant stage, the HY5-S36A-ox lines show strong growth inhibition (Supplemental Figure 5B, C). Taken together, our results clearly showed that higher binding affinity of unphosphorylated HY5 is physiologically more active than phosphorylated HY5 in regulating photomorphogenesis, especially during rapid light response.

**Figure 7.**
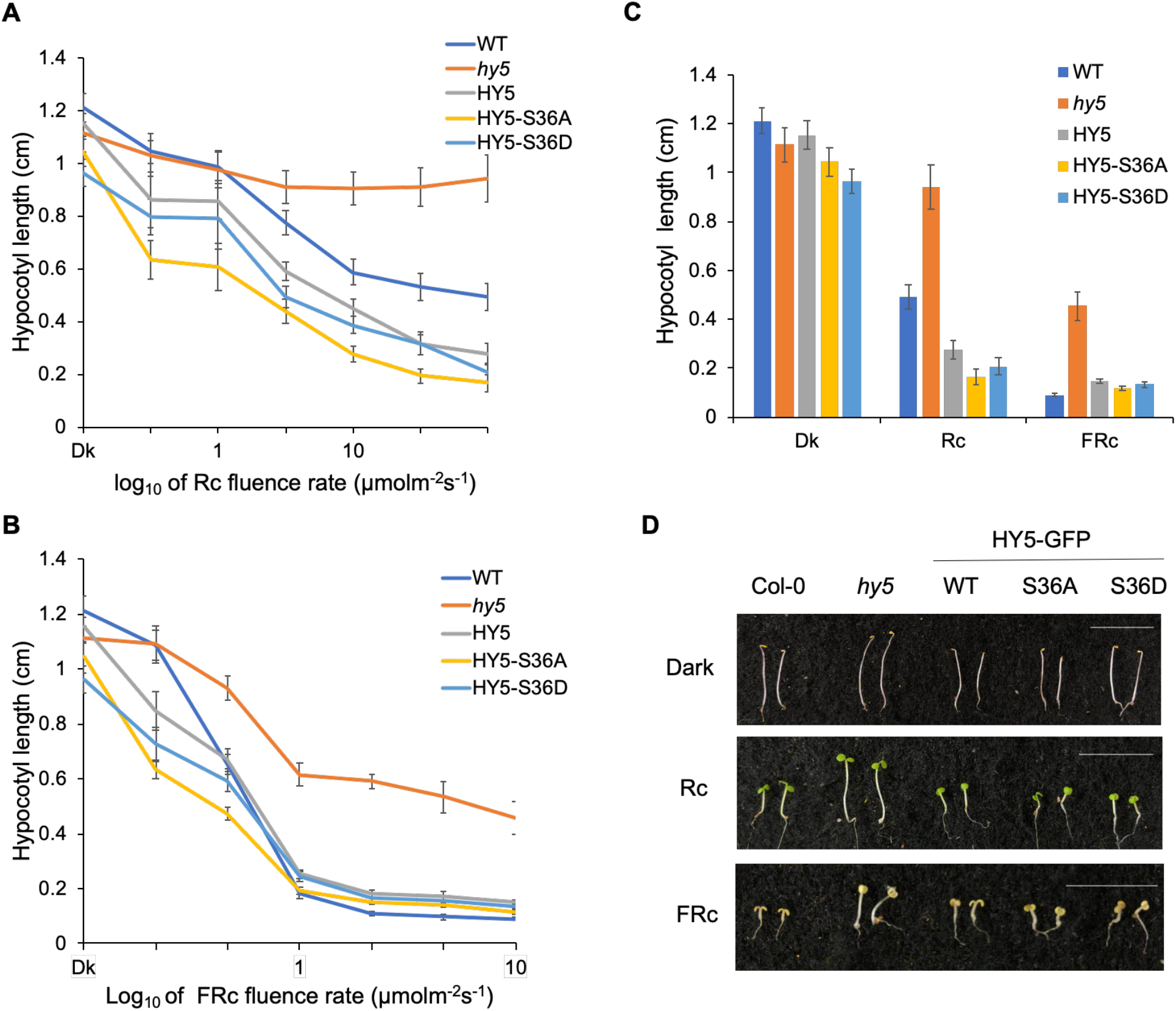
Unphosphorylated HY5 displays enhanced photomorphogenesis. **(A, B)** Line graphs show hypocotyl length of each HY5 overexpressing transgenic lines (HY5, HY5-S36A and HY5-S36D) in response to red light (Rc) **(A)** and far-red light (FRc) **(B)** with increasing light intensities. Error bars indicates SD (n > 30). **(C)** A bar graph shows the hypocotyl length of wild-type Col-0, *hy5* mutant, HY5, HY5-S36A and HY5-S36D overexpressing lines grown under dark (Dk), Red (Rc; 6 µmolm^-2^ s^-1^), and far-red light (FRc; 1 µmolm^-2^ s^-1^) conditions for 4 days. Error bars indicates SD (n > 30). **(D)** Photographs show the hypocotyl length of various genotypes in continuous Dark or Rc or FRc. Seeds of various genotypes were grown on MS medium without sucrose for 4 days. Scale bar: 10mm.

## DISCUSSION

HY5 protein plays a pivotal role in photomorphogenesis. Therefore, it is not surprising that a number of HY5 binding targets are involved in diverse developmental processes. In Arabidopsis, more than 60% of early-induced genes by phyA or phyB are HY5 direct targets, which strongly supports the notion that HY5 is one of the high hierarchical regulators of the transcriptional cascade for photomorphogenesis (Gangappa and Botto, 2016). The regulation of HY5 by light includes phytochromes, PIFs, and COP1-SPA E3 ubiquitin ligase complex. Previous studies have been focused on the roles of HY5 in the regulation of gene expression (Gangappa and Botto, 2016). Despite the phosphorylation in HY5 has been reported many years ago, the underlying mechanism and physiological roles have remained elusive. Here, we provide a mechanistic view on how HY5 is phosphorylated and ubiquitinated by a cognate kinase-E3 ubiquitin ligase (COP1-SPA) to modulate its regulatory activity in photomorphogenesis.

In this study, we provide evidence that SPA1 is the kinase for HY5 and thus regulates its stability and activity. We observed strong phosphorylation signals *in vitro* kinase assay and the phosphorylation of HY5 was significantly reduced in *spaQ* mutant compared with that in WT *in vivo* (Figures 1 and 2). We also found that SPA1 phosphorylates HY5 through only one phosphorylation site which is Ser-36 (Figure 1D). The mutation of this site abolished the phosphorylation of HY5. Previously, CKII was suspected to be a likely candidate kinase of HY5 (Hardtke *et al*., 2000). CKII was also considered as the kinase of PIFs and another positively acting bHLH transcription factor HFR1 (Park *et al*., 2008, Bu *et al*., 2011). However, the *in vitro* kinase assay using a kinase-containing fraction of enriched seedling extract in a previous study is not sufficient to conclude that CKII is the kinase given that other components may also exist in the extract. In conclusion, we presented strong evidence that SPA1 is the key kinase for HY5.

It has been suggested that the kinase domain of SPA1 may provide structural information critical for SPA1 function (Holtkotte *et al*., 2016). SPA1^R517E^ mutant shows a defect in PIF1 degradation and seed germination in response to light (Paik *et al*., 2019). Similarly, SPA1^R517E^ (mSPA1) transgenic lines failed to rescue the seedlings de-etiolation phenotype of *spaQ* (Supplemental Figure 2C). Since the Arg 517 in SPA1 is well conserved in many plants (Paik *et al*., 2019), it is possible that the structural integrity of the kinase domain is important for its proper function, while the biochemical basis remains unknown. Our study shows that the kinase domain of SPA1 may act as a molecular scaffold for potential protein-protein interaction. We observed considerable HY5 accumulation in the SPA1^R517E^ line (Supplemental Figure 2B), which explained the deetiolation phenotype of SPA1^R517E^ mutant, showing photomorphogenesis in the dark. Moreover, the accumulation of HY5 in SPA1^R517E^ mutant in the dark resulted from the failure of degradation by COP1-SPA complex confirmed our hypothesis that the kinase domain of SPA1 might act as a molecular scaffold for protein-protein interaction.

Phosphorylation of transcription factors is a common modification that can influence their biological properties such as multimerization or nucleocytoplasmic partitioning in both plants and animals. In plant photomorphogenesis, phosphorylation has been observed for many transcription factors (Pham *et al*., 2018). Other than PIF1, light can also induce phosphorylation of PIF3 at multiple sites, and phosphorylated PIF3 is subject to the degradation by LIGHT RESPONSIVE BTB PROTEIN (LRB) and EIN3-BINDING F BOX PROTEIN (EBF) E3 ligases (Ni *et al*., 2013, Ni *et al*., 2014). Phosphorylation can also regulate the transcriptional activity of PIF4 which affects diurnal hypocotyl elongation and also influence translocation of PIF7 which regulates shade-induced stem elongation (Bernardo-García *et al*., 2014, Dong *et al*., 2017, Huang *et al*., 2018). In this study, we demonstrated that the phosphorylation of HY5 results in lower binding affinity to COP1 and SPA1, which reduces HY5 ubiquitination and in turn stabilizes HY5 proteins in the dark (Figures 4 and 5). Therefore, HY5 abundance is regulated apparently by two parallel pathways. One is the oscillation of the availability of COP1 in the nucleus, and the other is the specific phosphorylation of HY5, which modulates HY5’s ability to interact with COP1-SPA complex. Considering that HY5 is a key positive regulator of photomorphogenesis at the early seedling stage, this mechanism may help maintain a small pool of less active HY5 in the dark so that seedlings can give a rapid initial response during dark-to-light transition. Thus, SPA1 is acting both negatively and positively to regulate HY5 level and activity, thus forming a negative feedback loop between HY5 and COP1-SPA. This is consistent with a recent study showing that accumulation of HY5 in the dark leads to an increase in the COP1/SPA complex, and thus its own degradation (Burko *et al*., 2020).

Phosphorylation of transcription factors to regulate their stability and DNA binding capacity is common in eukaryotic cells (Hunter, 2007). Previous ChIP-chip analysis showed that HY5 binds directly to the promoters of genes related to auxin signaling, ethylene signaling, and gibberellin signaling. Moreover, it was shown that HY5 is necessary for the rapid transcription of those genes during the dark-to-light transition, which eventually allows the accumulation of chlorophyll and anthocyanin for photosynthesis (Lee *et al*., 2007). We show here that unphosphorylated HY5 has a stronger binding affinity to its target promoters, such as G-box and ACE-box in *XTH15, EXP2, IAA19* and *SAUR36*, than phosphorylated HY5 (Figure 6A). This is consistent with the phenotype of HY5-S36A transgenic line which shows shorter hypocotyl length compared to wild type HY5 (Figure 7), which is also confirmed by qRT-PCR analyses (Figure 6B). HY5 activity is also regulated by interaction with other transcription factors including the BBX factors. In fact, HY5 lacks any transcriptional activation domain. A recent study showed that interaction with BBX20/21/22 proteins are necessary for activation of gene expression by HY5 (Bursch *et al*., 2020). Whether phosphorylation of HY5 has any influence on interaction with other transcription factors awaits further studies.

The phosphorylation of HY5 by SPAs reveals an intervention point for directed modulation of growth properties. Here we show that by modulating phosphorylation of HY5 and altering its protein stability and activity, plants can rapidly respond to light irradiation and also avoid over-photomorphogenesis, which would be of great advantage for seedlings in the constantly changing natural environment. A previous study reported that the kinase activity responsible for HY5 phosphorylation might be regulated by light (Hardtke *et al*., 2000), they proposed that unphosphorylated HY5 may accumulate more under the light for efficient light response. However, we observed phosphorylation of HY5 in both light and dark (Figure 2A). Whether any regulators acting as the rate-limiting factors of HY5 phosphorylation or SPA1 kinase activity needs to be further studied. We also show that unphosphorylated HY5 proteins might modulate growth in the post-seedling stage (Supplemental Figure 5B, C). It will be of interest to study how plants fine-tune some transcription factors by post-translational modification to induce optimal plant development.

## EXPERIMENTAL PROCEDURES

### Plant materials and growth conditions

Wild type Col-0, various mutants, and transgenic plants in the Col-0 background were used in this study, unless otherwise indicated. Plants were grown in soil under 24-h light at 22 ± 0.5 °C. TAP-SPA1, LUC-SPA1/*spaQ* and LUC-mSPA1/*spaQ* transgenic plants were reported previously (Paik *et al*., 2019). To generate HY5, HY5-S36A and HY5-S36D transgenic lines, the HY5 36th Serine (AGC) was changed into Alanine (GCC) and Aspartic Acid (GAC) respectively using Quickchange II site-directed mutagenesis kit (Agilent, Catalog #200523) and cloned into pB7FWG vectors, then transformed into *hy5* mutant, *spaQ* mutant and WT. The transformants were selected in the presence of Basta. Transgenic lines with same overexpressed HY5 proteins were used (Supplemental Figure 5A).

### Vector constructions and protein purification

MBP-COP1 and MBP-SPA1 were prepared as described previously (Xu *et al*., 2014, Paik *et al*., 2019). For generation of GST-HY5 and its mutant proteins, HY5, HY5-S36A and HY5-S36D were cloned into pGEX4T-1. Each plasmid was transformed into BL21(DE3) cells. Protein expression was induced under 16 °C for overnight with 0.1 mM IPTG. Collected cells were sonicated in binding buffer (100 mM Tris-Cl, pH 7.5, 150 mM NaCl, 0.2% Tergitol NP-40, 1x Protease inhibitor cocktail, and 1 mM PMSF) and purified using GST agarose beads (Pierce, Waltham, USA. Cat. # 20211). Proteins were eluted with the elution buffer (Tris-Cl, PH 7.5, 150 mM NaCl, 1 mM, 10mM glutathione, 10% Tergitol NP-40, 10% glycerol, 1mM PMSF, 1x Protease inhibitor cocktail) into separate fractions. The eluted proteins were analyzed on SDS-PAGE gel and used for kinase assays and pull-down assay.

### *In vitro* kinase assay

For SPA1 kinase assay, about 500 ng of MBP-SPA1 and 1 µg of GST-HY5 or GST-HY5-S36A fusion proteins were used. All kinase assays were performed in a kinase buffer (50 mM Tris, pH 7.5, 4 mM β-mercaptoethanol, 1 mM EDTA, 10 mM MgCl2). ^32^P radio-labeled gamma-ATP (Perkin Elmer Cat# BLU502A) was added to the reaction and incubated at 28 °C for 1 h, unless otherwise indicated. SDS sample buffer (6x) was added to stop the reaction and the boiled proteins were separated on SDS-PAGE gel. The gels were dried and exposed to a phosphor screen and then scanned with Typhoon FLA 9500 (GE healthcare, Chicago, USA).

### HY5 mobility shift assay

To observe GFP-HY5 mobility shift in Col-0 and *spaQ*, total protein was separated in a 10·10.5 cm 7% SDS-PAGE or 8% SDS-PAGE gel containing 20-50 µM phostag acrylamide (Wako Japan, Cat# AAL-107). Total proteins were extracted from 4-day-old light-grown or dark-grown seedlings with extraction buffer (100 mM Tris·HCl [pH 6.8], 20% glycerol, 5% SDS, 20 mM dithiothreitol, 1 mM PMSF, 1× protease inhibitor, and 100 µM bortezomib). The extracts were cleared by centrifugation and then incubated with or without 400 units/mL Alkaline Phosphatase, Calf Intestinal (CIP) (NEB cat # M0290) at 37 °C for 30min. The reaction mixtures were terminated by adding 6x SDS loading buffer and boiling at 99 °C for 10 min. Immunoblotting analyses were performed with anti-GFP and anti-Tubulin antibodies.

### Protein extraction and Western blot analyses

To analyze HY5 abundance in dark-to-light and light-to-dark transition, seeds were surface sterilized and grown in the dark or continuous light for four days or followed with according light or dark treatments. To analyze HY5 abundance in mutants and transgenic lines, seedlings were grown in continuous dark for four days. For total protein extraction, whole seedlings were collected and ground in 100 µL denaturing extraction buffer [100 mM Tris pH 7.5, 1 mM EDTA, 8 M Urea, 1x protease inhibitor cocktail (Sigma-Aldrich Co., St. Louis, MO, cat# 59), 2 mM PMSF] and cleared by centrifugation at 20,200 × g for 10 min at 4 °C. Samples were boiled for 10 min with 6X SDS sample buffer and separated on a 10% SDS-PAGE gel, blotted onto PVDF membranes, and probed with corresponding antibodies. Antibodies used in these studies are anti-HY5 (Abiocode, Cat. # R1245-2), anti-GFP (Abcam, Cat. # ab6556 for immunoprecipitation), anti-Tubulin antibodies (Enzo Life Sciences, Cat. # BML-PW8770-0025). Secondary HRP bound antibodies were visualized with Super Signal West Pico Chemiluminescent substrate (Pierce Biotechnology Inc., Waltham, USA.), and developed with an X-ray film or GBox-F3 Syngene Imager. The HY5 amount was quantified with image J from three independent blots and normalized by its corresponding Tubulin levels.

### *In vitro* pull-down assay

For *in vitro* pull-down assays, MBP-COP1, MBP-SPA1 and GST-HY5, GST-HY5-S36A and GST-HY5-S36D fusion proteins were prepared as described previously. 1ug protein was used for each of them. All protein combinations were incubated with 20 mL of amylose resin in the binding buffer (50 mM Tris-Cl, pH 7.5, 150 mM NaCl, 0.6% Tween 20, and 1 mM DTT) for 3 h. The beads were collected and washed six times with 5 min of rotation each time in binding buffer. The bound HY5 was detected by anti-GST-HRP conjugate (RPN1236; GE Healthcare Bio-Sciences). Membranes were developed and visualized as described above. The intensity of the GST-HY5 band from three independent blots was quantified using ImageJ software. These relative values are shown as bar graphs.

### *In vivo* co-IP assay

For Co-IP experiments, homozygous HY5-GFP, HY5-S36A and HY5-S36D transgenic seedlings were grown in dark for 4 days and then treated with 40 µM bortezomib (LC Laboratories, Woburn, MA) for at least 4h. Total proteins were extracted from 1g tissue with 1ml protein extraction buffer. After 15 min centrifugation at 16,000g at 4°C in darkness, 100 ul supernatant of each sample was reserved as total, and the remainder was incubated with Dynabeads Protein A (Life Technologies Co., Carlsbad, CA, Cat. No: 10002D) bound with anti-GFP antibody (Abcam, Cambridge, MA, Cat. No: ab9110). Twenty μl Dynabeads with 1g antibody were used for individual sample. After 2 h incubation in the dark at 4°C, beads were washed three times with 1 ml protein extraction buffer with 0.2% NP40. Immuno-precipitated proteins were analyzed by immunoblotting.

### RNA extraction and quantitative RT-PCR

The quantitative RT-PCR (qRT-PCR) analysis was performed as described with minor variations (Shor *et al*., 2017). Total RNA was isolated from 4-day-old dark-grown seedlings plus 3hrs light treatment using the Spectrum Plant Total RNA Kit (Sigma-Aldrich). One microgram of total RNA was treated with DNase I to eliminate genomic DNA and then reverse transcribed using SuperScript III (Life Technologies) as per the manufacturer’s protocol. Real-time PCR was performed using the Power SYBR Green RT-PCR Reagents Kit (Applied Biosystems) in a 7900HT Fast Real-Time PCR machine (Applied Biosystems). *PP2A* was used as a control to normalize the expression data. The resulting cycle threshold values were used to calculate the levels of expression of different genes relative to PP2A, as suggested by the manufacturer (Applied Biosystems). Primer sequences used for qRT-PCR are listed in Supplemental Table 1.

### ChIP-qPCR assay

Three biological replicates of HY5 and HY5-S36A and HY5-S36D seedlings grown in the dark for 4 days and moved to simulated white light for 3hrs were used for ChIP-qPCR analysis. ChIP experiment was performed as previously described (Shor *et al*., 2017). Anti-GFP (Abcam, Cat. # ab6556 for immunoprecipitation) antibody was used for immunoprecipitation. After elution, reversing crosslinks, and DNA purification, the amount of each precipitated DNA fragment was detected by real-time qPCR using specific primers listed in Supplemental Table 1. Three biological replicates were performed, and three technical repeats were carried out for each biological replicate. Representative results from one biological replicate were shown.

### Measurement of hypocotyl lengths

For the measurement of hypocotyl length under dark, red light (Rc) and far-red light (FRc) with different intensities, images of 150 seedlings (30 seedlings for each line with three independent biological replicates) at each light intensity condition were taken and then measured using the publicly available IMAGEJ software (http://rsb.info.nih.gov/ij/). Seeds were plated on MS medium without sugar and kept in the dark for 3 days at 4°C. Seeds were then exposed to 3 hrs of white light to induce germination and then kept in the dark for 21hrs. The dark grown plate seedlings were exposed in far-red light for additional 10min before being put in darkness. All the other plates were then put in the according conditions for 3 days as indicated in the figure. Light fluence rates were measured using a spectroradiometer (Model EPP2000; StellarNet) as described previously (Shen *et al*., 2005).

## ACKNOWLEDGMENTS

We thank Drs. Xing Wang Deng for sharing the *cop1* mutants, and Ute Hoecker for sharing the *spaQ* mutant. We thank the Huq and Sung lab members for critical reading of the manuscript. This work was supported by the grants from the National Science Foundation (MCB-2014408 to E.H. and IOS-1656764 to S.S.) and National Institute of Health (NIH) (GM-114297 to E.H. and GM-100108 to S.S.) The funders had no role in study design, data collection and analysis, decision to publish, or preparation of the manuscript.

## AUTHOR CONTRIBUTIONS

W.W, I.P., J.K., S.S. and E.H. designed experiments. W.W, I.P. and J.K. carried out experiments. W.W, I.P., J.K., S.S. and E.H. analyzed data. W.W, and J.K. wrote the article. X.H., S.S. and E.H. commented on the manuscript.

## SUPPORTING INFORMATION

Additional Supporting Information may be found in the online version of this article.

**Supplemental Figure 1:** Phosphorylation of HY5 occurs in both dark and light conditions.

**Supplemental Figure 2:** SPA1 kinase domain is necessary for its kinase activity and molecular function.

**Supplemental Figure 3:** Accumulation of HY5 protein in different genotypes.

**Supplemental Figure 4:** Co-immunoprecipitation (Co-IP) assays showing that non-phosphorylated form of HY5 proteins strongly interact with COP1 protein *in vivo*.

**Supplemental Figure 5:** Unphosphorylated HY5 lines show reduced growth.

**Supplemental Table 1:** Primer sequences used in experiments described in the text.

## Supplementary Information

**Supplemental Figure 1.**
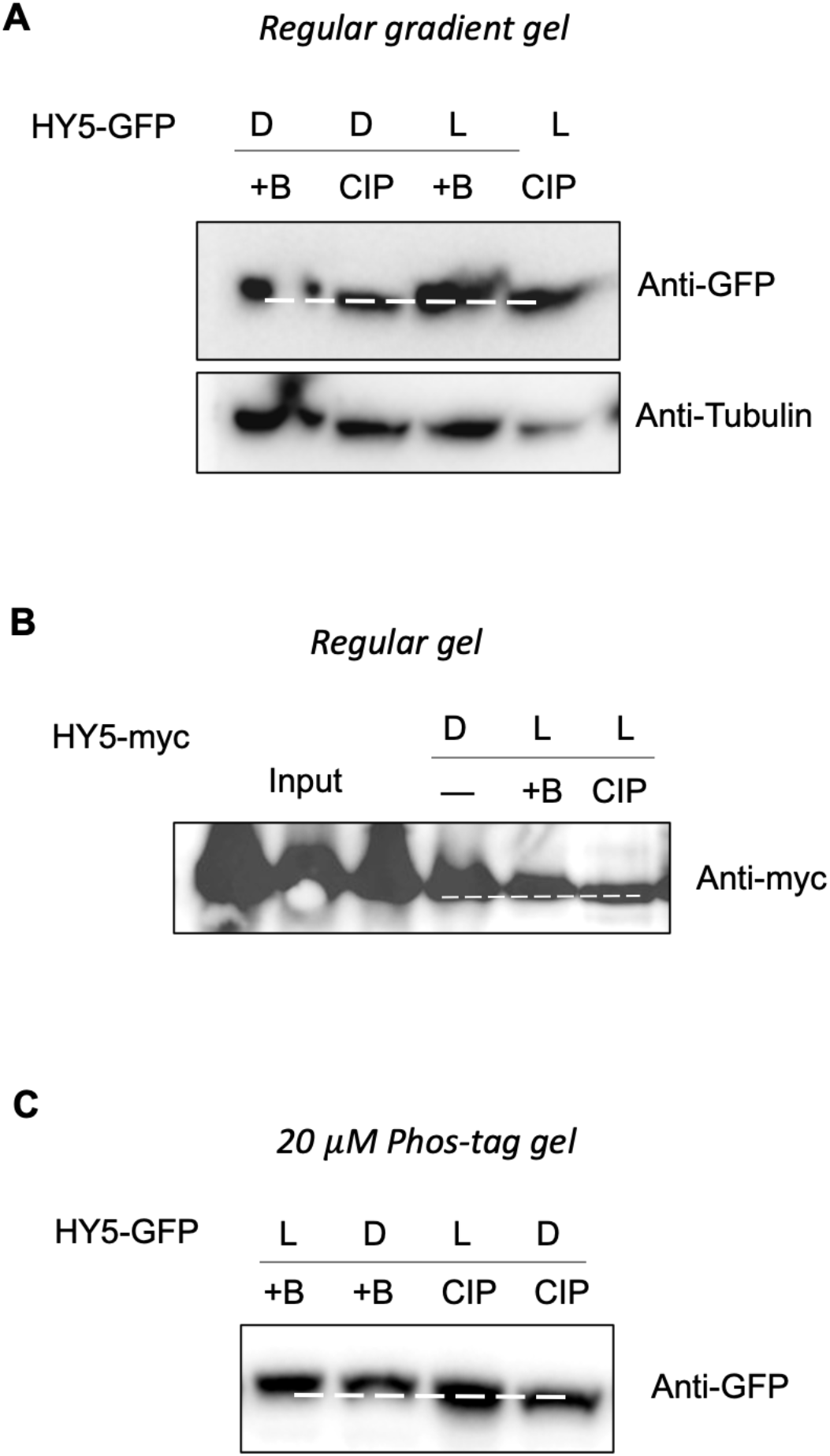
Phosphorylation of HY5 occurs in both dark and light conditions. **(A)** Immunoblots showing phosphorylation of HY5-GFP under both dark (D) and light (L) conditions. **(B)** Immunoblots showing phosphorylation of HY5-myc under light condition. **(C)** Immunoblots showing phosphorylation of HY5-GFP under both dark (D) and light (L) conditions in gels containing 20 µM phos-tag. Seedlings were grown in either dark or continuous light for 4 days before sampling for protein extraction. Protein was immunoprecipitated from transgenic plants and separated by SDS-PAGE (with or without phos-tag reagent) and tested with an anti-GFP or anti-myc antibody. The slow-migrating band is a phosphorylated form of HY5 as indicated by the phosphotase treatment. CIP, Calf Intestinal Alkaline Phosphatase; +B, inactivated boiled CIP; -, without CIP treatment.

**Supplemental Figure 2.**
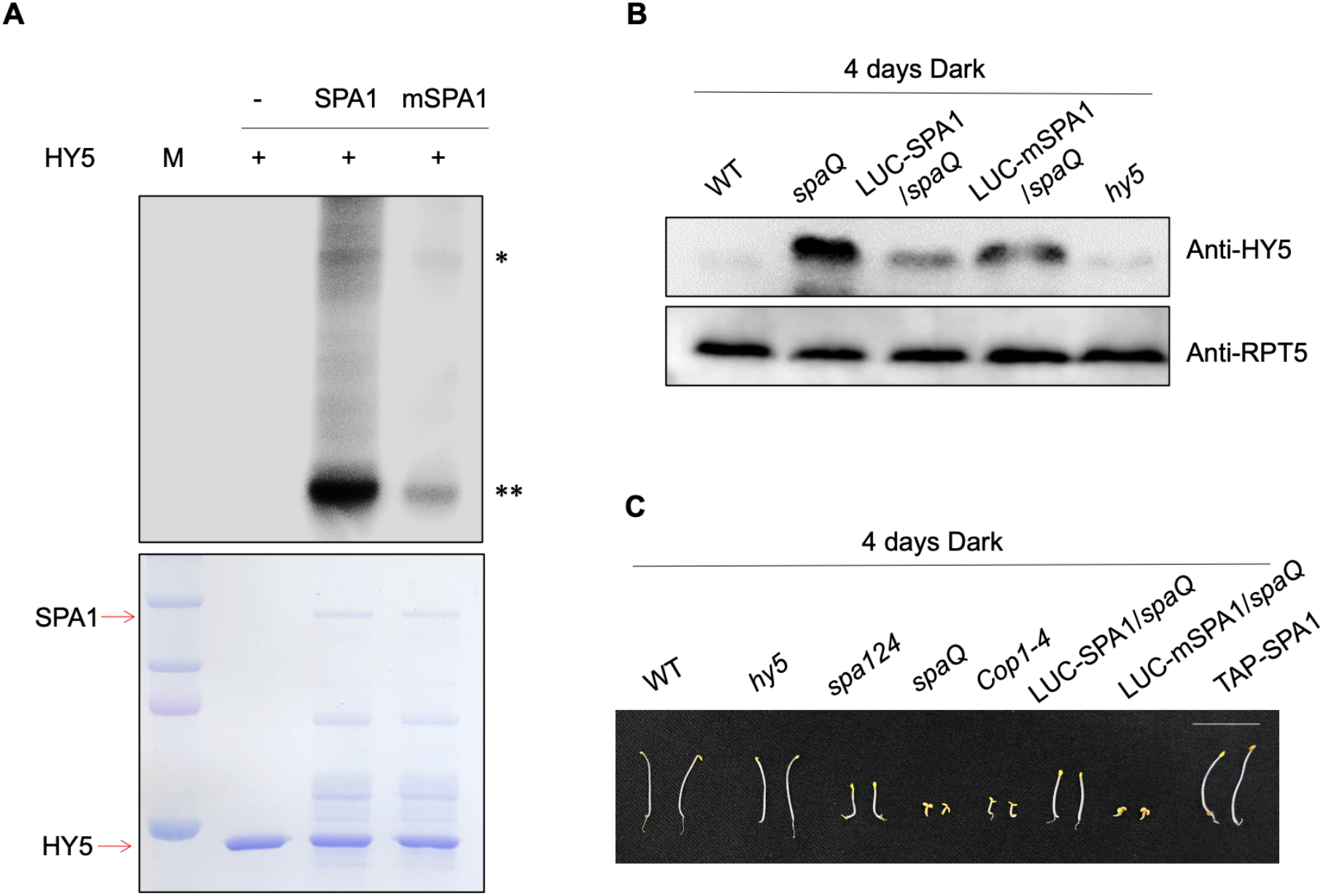
SPA1 kinase domain is necessary for its kinase activity and molecular function. **(A)** A conserved amino acid mutation on the SPA1 kinase domain (mSPA1) reduces the phosphorylation activity of SPA1 on HY5 (autoradiogram on top panel). *In vitro* kinase assay was performed using purified SPA1 and mSPA1 proteins from *Pichia pastoris* and GST-HY5 proteins from *E. coli*. The bottom panel shows the protein level in a Coomassie- stained gel. Single asterisk mark (*) shows SPA1 band and double asterisk mark (**) shows HY5 band. **(B)** Immunoblots showing HY5 accumulation in the different genotypes. Seedlings were grown in the dark for 4 days before sampling for protein extraction, endogenous anti-HY5 antibody and anti-RPT5 antibody were used. RPT5 protein was used as loading control. **(C)** Photograph showing the seedling phenotypes of different genotypes grown in darkness for 4 days. Scale bar: 10mm.

**Supplemental Figure 3.**
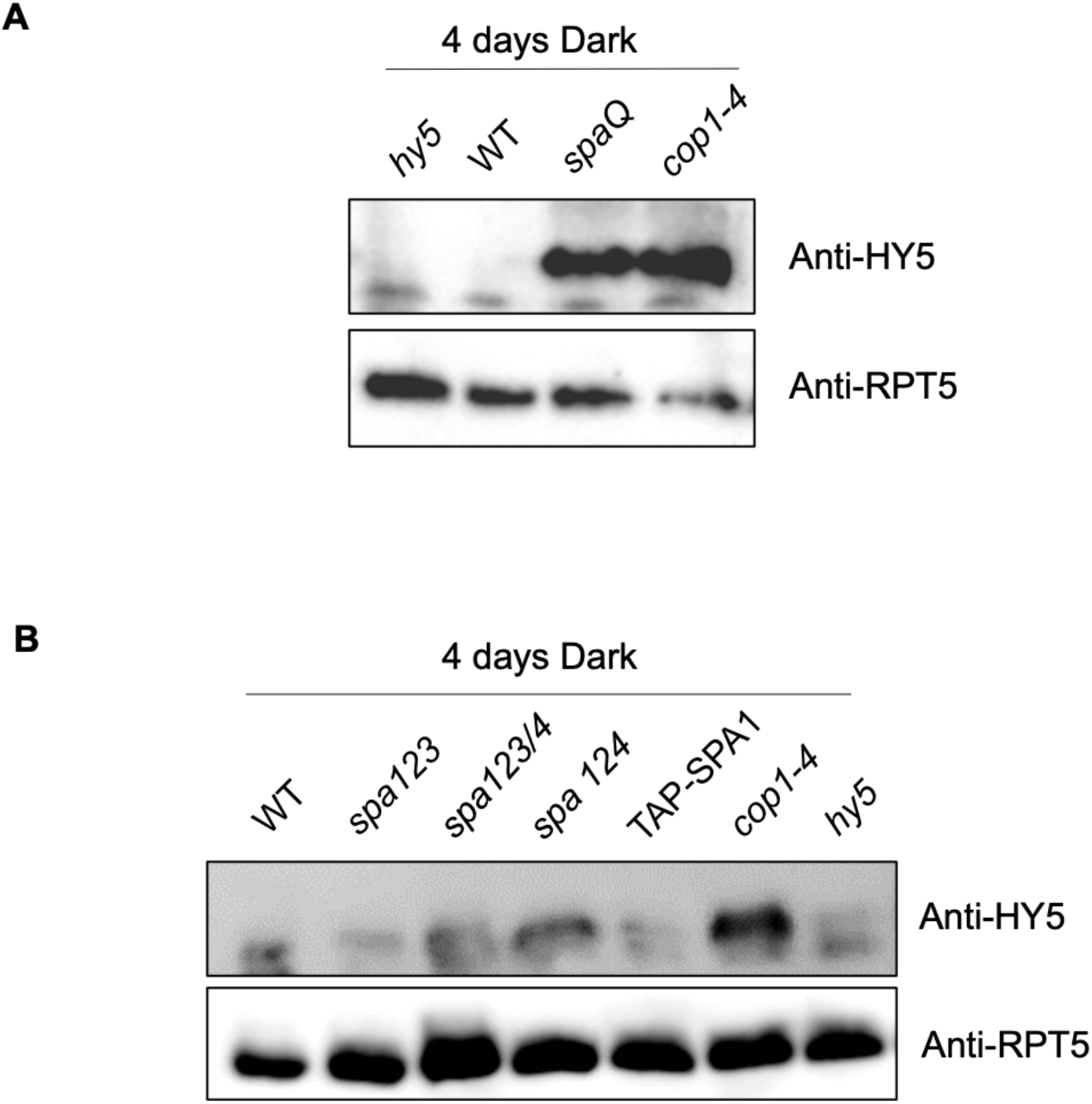
Accumulation of HY5 protein in different genotypes. **(A and B)** Seedlings were grown in the dark for 4 days before sampling for protein extraction. Endogenic HY5 protein level was detected by endogenous anti-HY5 antibody, RPT5 was used as loading control.

**Supplemental Figure 4.**
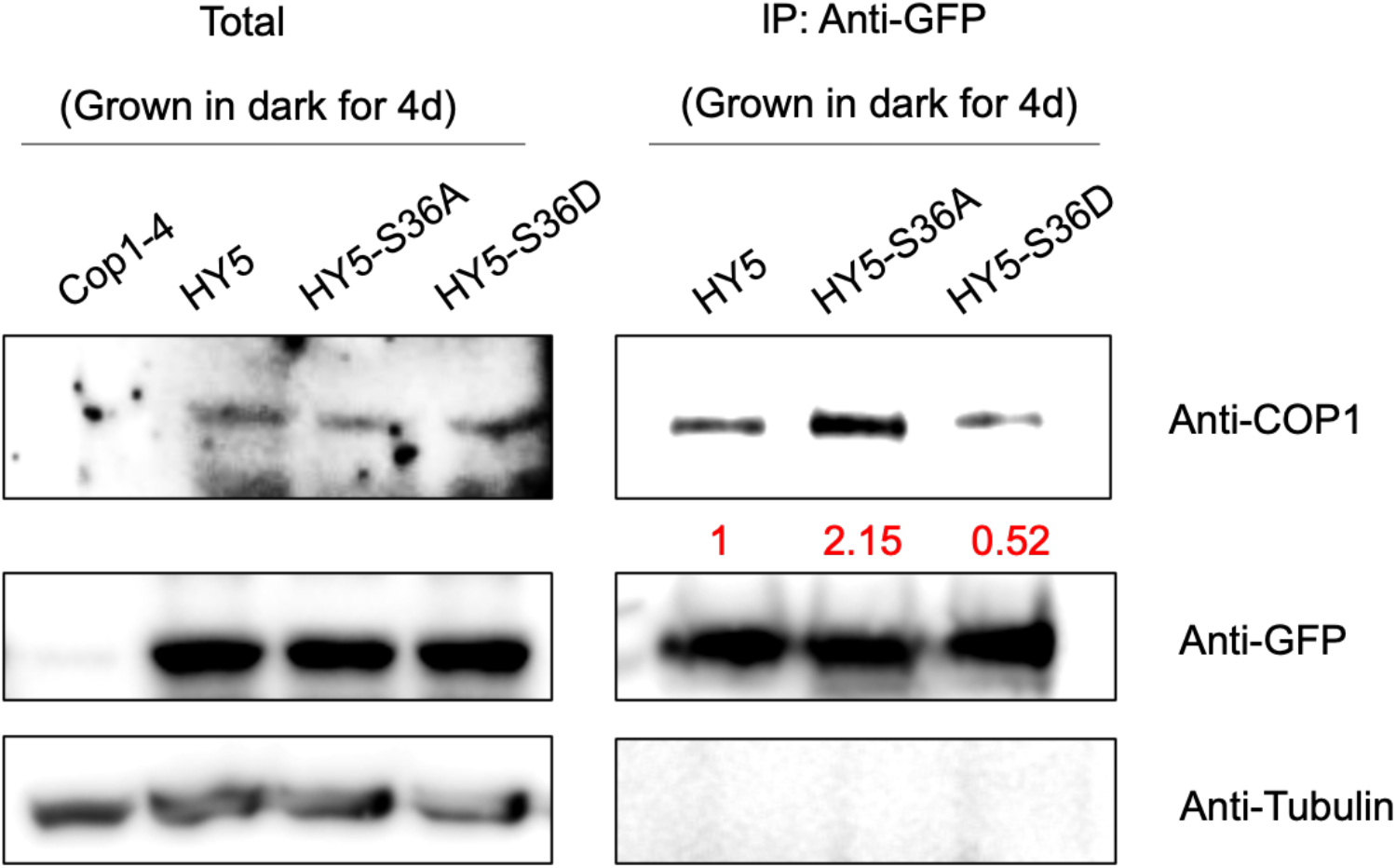
Co-immunoprecipitation (Co-IP) assays showing that non-phosphorylated forms of HY5 proteins strongly interact with COP1 protein *in vivo*. Homozygous HY5, HY5-S36A and HY5-S36D-GFP seedlings were grown at 22°C in continuous dark for 4 days, and then treated with 40 µM bortezomib for at least 4h. The total proteins were extracted and incubated with protein A beads. The total and precipitated proteins were examined by immunoblotting using antibodies against COP1, GFP and Tubulin, respectively. *cop1-4* mutant was used as the negative control. The numbers below anti-COP1 blots indicate the relative band intensities of co-precipitated COP1 normalized to those of precipitated HY5-GFP, HY5-S36A and HY5-S36D, respectively. The ratio of the first clear band was set to 1 for each blot.

**Supplemental Figure 5.**
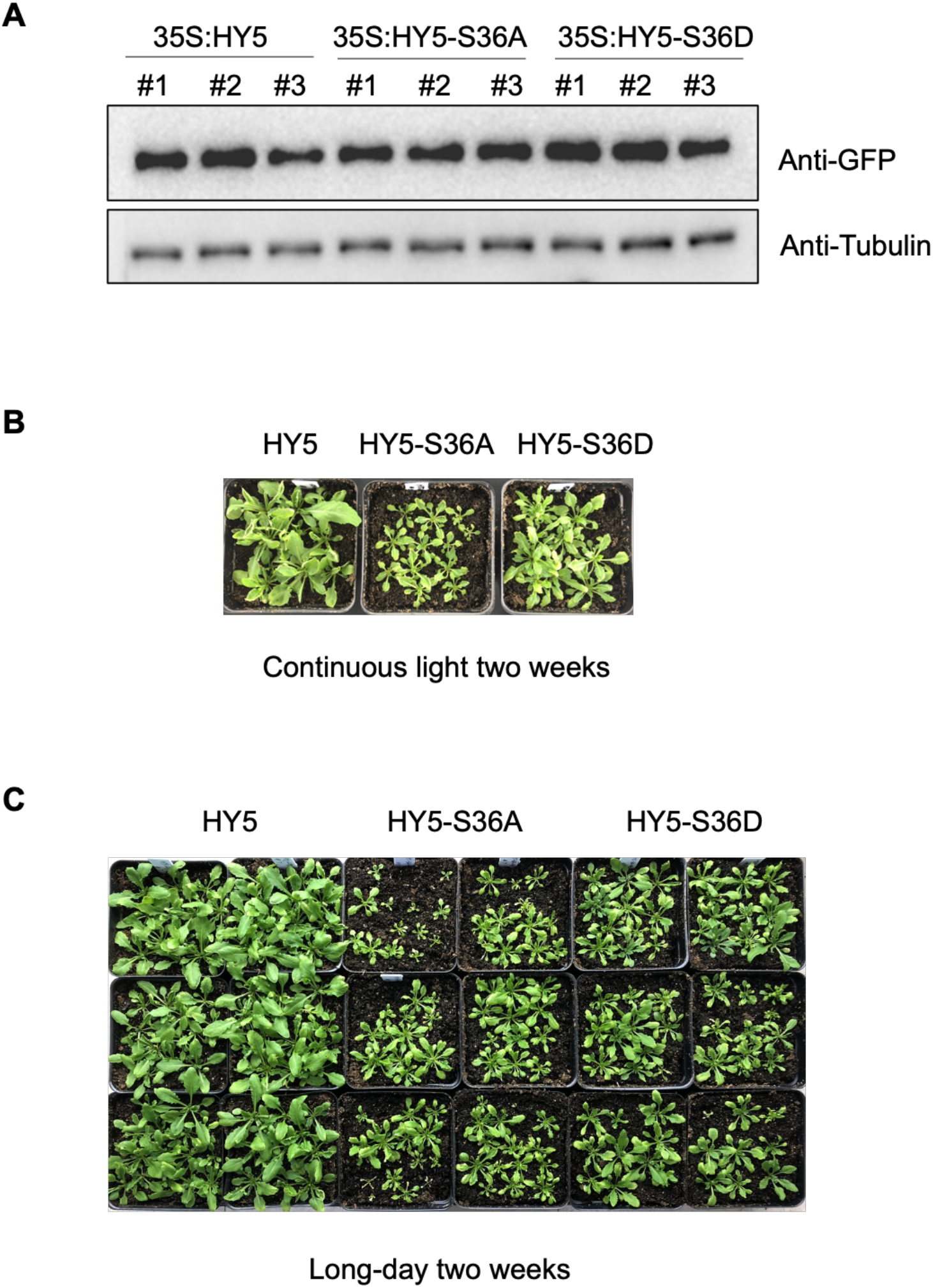
Unphosphorylated HY5 lines show reduced growth. **(A)** HY5 protein levels in three transgenic lines (HY5, HY5-S36A and HY5-S36D-GFP) used in this study. Proteins were extracted from 4 days light grown seedlings. **(B and C)** Adult phenotypes of three transgenic lines (HY5, HY5-S36A and HY5-S36D) grown in continuous light condition for two weeks **(B)** or in long-day condition for two weeks **(C)**.

**Supplemental Table 1.**
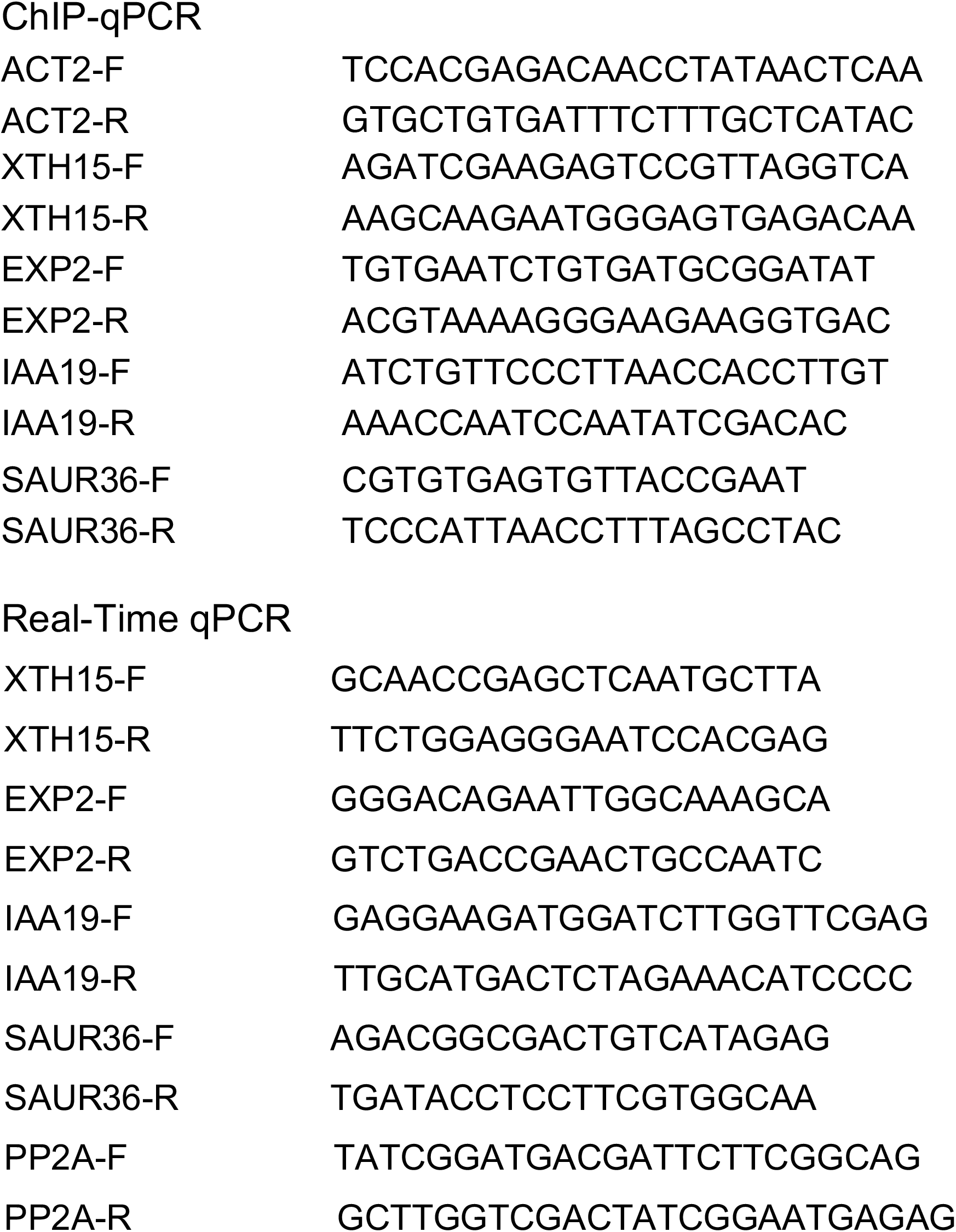
Oligo primers used in this study

