## Supplemental information for "Direct phosphorylation of HY5 by SPA1 kinase to regulate photomorphogenesis in Arabidopsis"

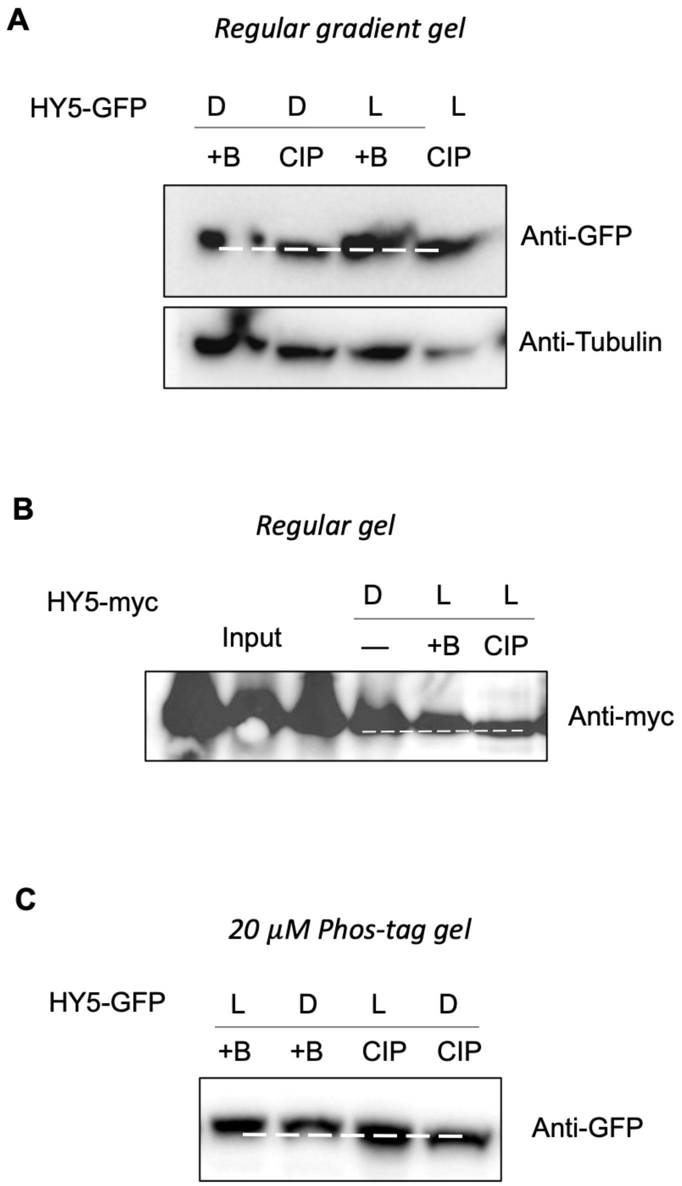


**Supplemental Figure 1. Phosphorylation of HY5 occurs in both dark and light conditions.** **(A)** Immunoblots showing phosphorylation of HY5-GFP under both dark (D) and light (L) conditions. **(B)** Immunoblots showing phosphorylation of HY5-myc under light condition. **(C)** Immunoblots showing phosphorylation of HY5-GFP under both dark (D) and light (L) conditions in gels containing 20 µM phos-tag. Seedlings were grown in either dark or continuous light for 4 days before sampling for protein extraction. Protein was immunoprecipitated from transgenic plants and separated by SDS-PAGE (with or without phos-tag reagent) and tested with an anti-GFP or anti-myc antibody. The slow-migrating band is a phosphorylated form of HY5 as indicated by the phosphotase treatment. CIP, Calf Intestinal Alkaline Phosphatase; +B, inactivated boiled CIP; -, without CIP treatment.

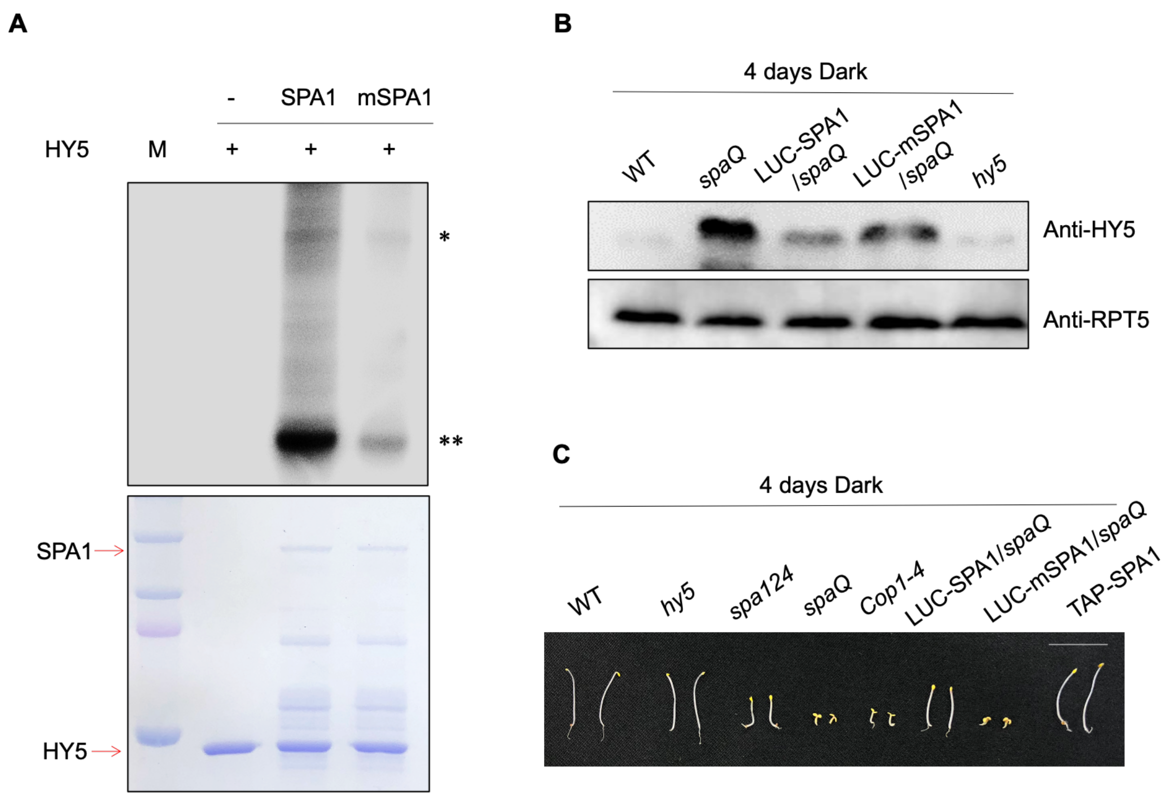


**Supplemental Figure 2. SPA1 kinase domain is necessary for its kinase activity and molecular function.**

**(A)** A conserved amino acid mutation on the SPA1 kinase domain (mSPA1) reduces the phosphorylation activity of SPA1 on HY5 (autoradiogram on top panel). *In vitro* kinase assay was performed using purified SPA1 and mSPA1 proteins from *Pichia pastoris* and GST-HY5 proteins from *E. coli*. The bottom panel shows the protein level in a Coomassie- stained gel. Single asterisk mark (*) shows SPA1 band and double asterisk mark (**) shows HY5 band. **(B)** Immunoblots showing HY5 accumulation in the different genotypes. Seedlings were grown in the dark for 4 days before sampling for protein extraction, endogenous anti-HY5 antibody and anti-RPT5 antibody were used. RPT5 protein was used as loading control. **(C)** Photograph showing the seedling phenotypes of different genotypes grown in darkness for 4 days. Scale bar: 10mm.

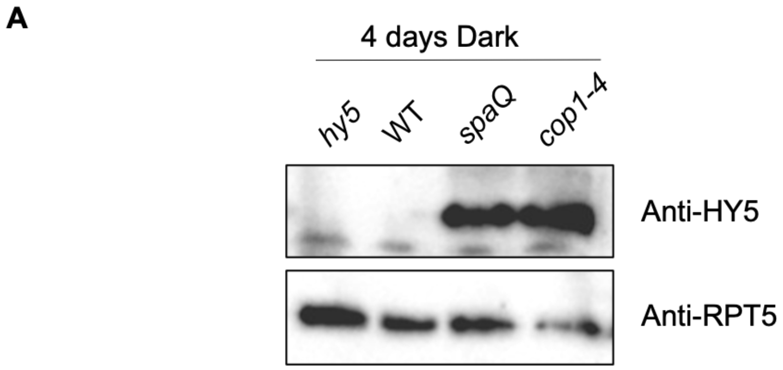


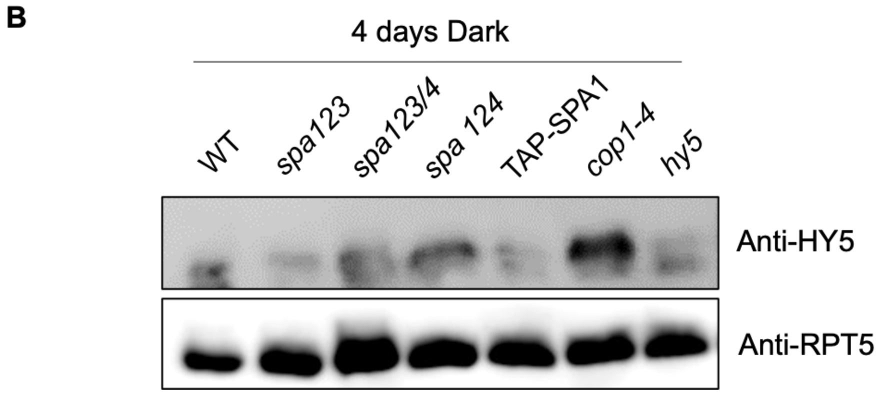


**Supplemental Figure 3. Accumulation of HY5 protein in different genotypes.**
**(A and B)** Seedlings were grown in the dark for 4 days before sampling for protein extraction. Endogenic HY5 protein level was detected by endogenous anti-HY5 antibody, RPT5 was used as loading control.

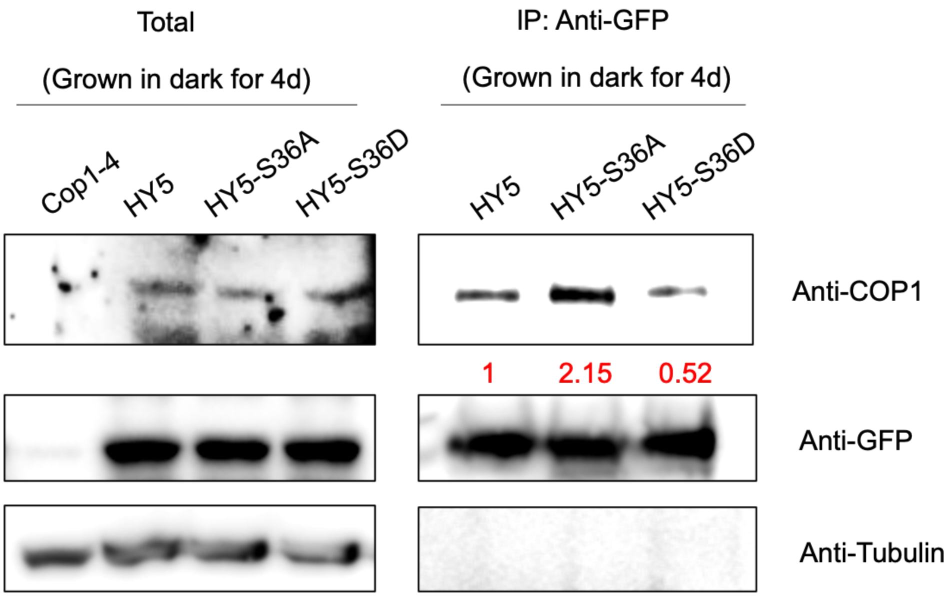


**Supplemental Figure 4. Co-immunoprecipitation (Co-IP) assays showing that non-phosphorylated forms of HY5 proteins strongly interact with COP1 protein *in vivo*.**
Homozygous HY5, HY5-S36A and HY5-S36D-GFP seedlings were grown at 22°C in continuous dark for 4 days, and then treated with 40 μM bortezomib for at least 4h. The total proteins were extracted and incubated with protein A beads. The total and precipitated proteins were examined by immunoblotting using antibodies against COP1, GFP and Tubulin, respectively. *cop1-4* mutant was used as the negative control. The numbers below anti-COP1 blots indicate the relative band intensities of co-precipitated COP1 normalized to those of precipitated HY5-GFP, HY5-S36A and HY5-S36D, respectively. The ratio of the first clear band was set to 1 for each blot.

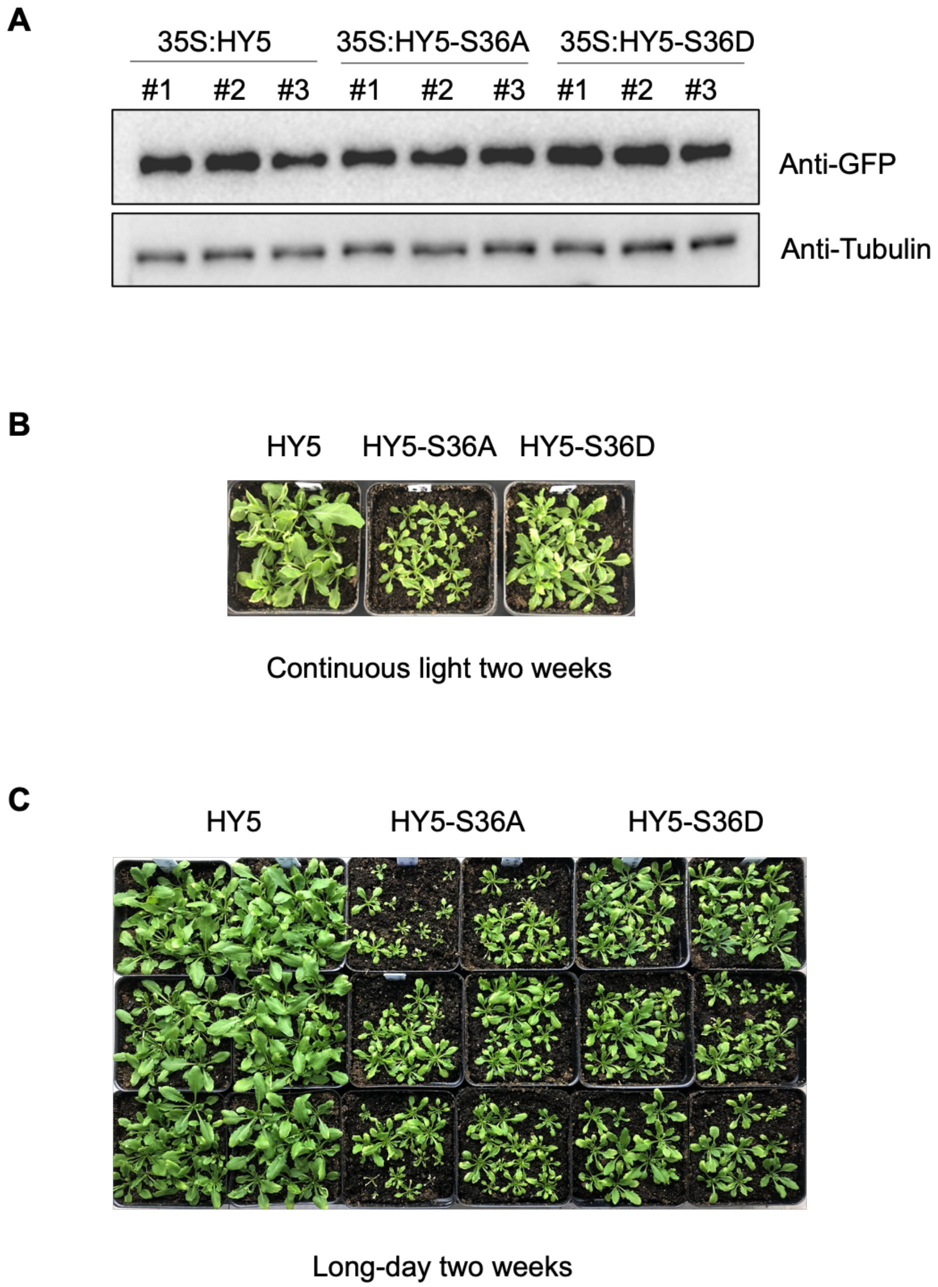


**Supplemental Figure 5. Unphosphorylated HY5 lines show reduced growth.**
**(A)** HY5 protein levels in three transgenic lines (HY5, HY5-S36A and HY5-S36D-GFP) used in this study. Proteins were extracted from 4 days light grown seedlings. **(B and C)** Adult phenotypes of three transgenic lines (HY5, HY5-S36A and HY5-S36D) grown in continuous light condition for two weeks **(B)** or in long-day condition for two weeks **(C)**.

**Supplemental Table 1.** Oligo primers used in this study

| | |
|---|---|
| **ChIP-qPCR** | |
| ACT2-F | TCCACGAGACAACCTATAACTCAA |
| ACT2-R | GTGCTGTGATTTCTTTGCTCATAC |
| XTH15-F | AGATCGAAGAGTCCGTTAGGTCA |
| XTH15-R | AAGCAAGAATGGGAGTGAGACAA |
| EXP2-F | TGTGAATCTGTGATGCGGATAT |
| EXP2-R | ACGTAAAAGGGAAGAAGGTGAC |
| IAA19-F | ATCTGTTCCCTTAACCACCTTGT |
| IAA19-R | AAACCAATCCAATATCGACAC |
| SAUR36-F | CGTGTGAGTGTTACCGAAT |
| SAUR36-R | TCCCATTAACCTTTAGCCTAC |
| **Real-Time qPCR** | |
| XTH15-F | GCAACCGAGCTCAATGCTTA |
| XTH15-R | TTCTGGAGGGAATCCACGAG |
| EXP2-F | GGGACAGAATTGGCAAAGCA |
| EXP2-R | GTCTGACCGAACTGCCAATC |
| IAA19-F | GAGGAAGATGGATCTTGGTTCGAG |
| IAA19-R | TTGCATGACTCTAGAAACATCCCC |
| SAUR36-F | AGACGGCGACTGTCATAGAG |
| SAUR36-R | TGATACCTCCTTCGTGGCAA |
| PP2A-F | TATCGGATGACGATTCTTCGGCAG |
| PP2A-R | GCTTGGTCGACTATCGGAATGAGAG |
